# DNA-enhanced CuAAC ligand enables live-cell detection of intracellular biomolecules

**DOI:** 10.1101/2022.11.10.515969

**Authors:** Keqing Nian, Yifang Liu, Yuchen Qiu, Zhuoyu Zhang, Laura Brigandi, Meni Wanunu, Sara H. Rouhanifard

## Abstract

Of the various conjugation strategies for cellular biomolecules, Cu(I)-catalyzed azide-alkyne cycloaddition (CuAAC) is the preferred click chemistry approach due to its fast reaction rate and the commercial availability of a wide range of conjugates. While extracellular labeling of biomolecules using CuAAC has been widely adopted, intracellular labeling in live cells has been challenging as the high copper concentrations required for CuAAC reaction is toxic to biological systems. As a critical first step towards CuAAC-mediated intracellular labeling, an ultrasensitive CuAAC ligand is needed to reduce cytosolic copper concentrations while maintaining fast reaction kinetics. Here, we developed BTT-DNA, a new DNA oligomer-conjugated CuAAC ligand for click reaction biomolecular labeling. The DNA oligo attachment serves several purposes, including: 1. Increased localization of copper atoms near the ligand, which enables ligation of azide tags with much lower copper concentrations than commercially available CuAAC ligands and without the addition of exogenous copper salt; 2. Allows nucleic acid template-driven proximity ligation by choosing the attached DNA sequence, 3. Enables the liposome encapsulation and delivery of the ligand into live cells, and 4. Facilitates intracellular labeling of nascent phospholipids in live cells. We demonstrate that BTT-DNA mediated labeling has little to no effect on the overall cell health.

## Introduction

Bioorthogonal chemistry comprises a set of broadly applied conjugation strategies used to study biological processes^1–4^. Among its primary requirements are a high degree of chemical specificity, rapid reaction times, and no side reactivity with native functional groups^5^. One of the first bioorthogonal reactions reported was Cu(I)-catalyzed azide-alkyne cycloaddition (CuAAC), whereby a dipolar cycloaddition between a chemically inert terminal alkyne and an azide is facilitated ^6–10^. The CuAAC reaction is compatible with aqueous conditions^11^, is usually free of byproducts^1^, and benefits from the commercial availability of a wide range of conjugates. However, the high copper catalyst concentration required in combination with ascorbate and oxygen can result in reactive oxygen species (ROS) that can be detrimental to nucleic acids^12^ and toxic to biological systems^13^. This toxic effect limits the utility of the CuAAC reaction inside live cells.

Biocompatible ligands have been developed to reduce the toxicity of the copper catalyst and enhance the reaction kinetics ^8,9^, for example, copper-chelating azides, which reduce the total copper requirement for the reaction^14^. However, even with these developments, few studies have achieved intracellular labeling using CuAAC in live cells due to copper toxicity at the necessary concentration range (typically 50-100 µM), low uptake of reagents, and competing intracellular ligands capable of sequestering the copper. For example, a copper-chelating azide was used to enhance CuAAC reactivity^15^ to detect an alkyne-labeled drug; however, in this strategy, the cells are fixed before fluorescent labeling. In another example, the CuAAC accelerating ligand was conjugated to a cell-penetrating peptide and demonstrated to detect l-homopropargylglycine^16^. However, the cell viability is diminished, with only 75% of cells viable after a 10-minute reaction due to the copper toxicity, and the intracellular click efficiency for cytosolic proteins is very low, with only 0.8% product yield after a 10-minute reaction. Therefore, CuAAC in live cells is typically used to label extracellular biomolecules on the cell surface such as glycans^8^ and lipids^17^. Alternatively, an azide and alkyne reporters may be introduced to live cells, however, cell fixation is performed before CuAAC-driven ligation of detection probes ^18,19^.

Copper-free, strain-promoted azide-alkyne cycloaddition (SPAAC) may also be used for covalent tagging of biomolecules^20^. This method is highly selective ^21,22^ and may be used for intracellularly labeling biomolecules^23^. However, SPAAC is limited by the commercial availability of cyclooctynes ^24^, reagent instability, and nucleophilic addition with cellular nucleophiles^20^. The inverse electron demand Diels-Alder reaction (IEDDA) may be used for intracellular labeling in live cells^25^ by using tetrazine-dye conjugates, which can show strong fluorescence intensity due to the reduction of tetrazine quenching interaction on the dye upon reaction with alkene However, tetrazine-alkene ligation is limited by the instability of tetrazine in aqueous solution^26^, and addition of the non-natural dienophiles, and the availability of activation methods^27^. Thus, developing a CuAAC reaction that may be used for intracellular, live-cell labeling is desirable.

We have developed here BTT-DNA, a new DNA oligomer-conjugated CuAAC accelerating ligand that enables the CuAAC reaction without adding exogenous copper salt. We show that the DNA oligo attachment localizes copper ions near the ligand, thus promoting CuAAC-mediated azide tagging with much lower copper concentrations than commercially available CuAAC ligands. The DNA oligo attachment also enabled a nucleic acid template-driven proximity ligation of a fluorogenic dye. We applied this proximity ligation to detect individual RNA molecules in the cell nucleus and cytoplasm using click RNA FISH assisted by our DNA oligomer-conjugated CuAAC ligand in fixed cells. DNA conjugation also provided a convenient method for intracellular delivery to encapsulate the copper within a liposome. We demonstrated that BTT-DNA protects the live cells from the toxicity and cellular perturbation caused by Cu(I) and sodium ascorbate and enables the live-cell, intracellular labeling of nascent phospholipids. In short, BTT-DNA allows sensitive detection of biomolecules in fixed and live cells, advancing our efforts toward applying CuAAC for intracellular, live-cell applications.

## Results

### Synthesis and characterization of BTT-DNA CuAAC accelerating ligand

We adopted the strategy used to develop the BTTAA ligand^9^ by ligating the precursor molecule (S1) that contains two *tert*-butyl groups bearing a single alkyne to a single-stranded DNA oligonucleotide probe bearing a single 3’ azide via CuAAC to produce BTT-DNA (**Fig. 1a**). We confirmed the synthesis of this probe by observing a gel shift following conjugation (**Supplementary Fig. 1**), then purified the BTT-DNA ligand with dialysis. Our early CuAAC studies using the BTT-DNA ligand suggested forming a ligation product without free copper (**Supplementary Fig. 1**). Using inductively coupled plasma mass spectrometry (ICP-MS), we assessed the presence of copper in the purified ligand, and found that the ratio of Cu: BTT-DNA is 10:1 (**Fig. 1a**). Interestingly, this Cu remained bound after several rounds of dialysis and HPLC purification. The same azide-modified DNA oligonucleotide was subjected to the CuAAC reaction conditions without S1 and assessed for copper. This produced a ratio of Cu: DNA of 7:1, indicating that the DNA and the triazole groups are chelating the copper.

**Fig 1.**
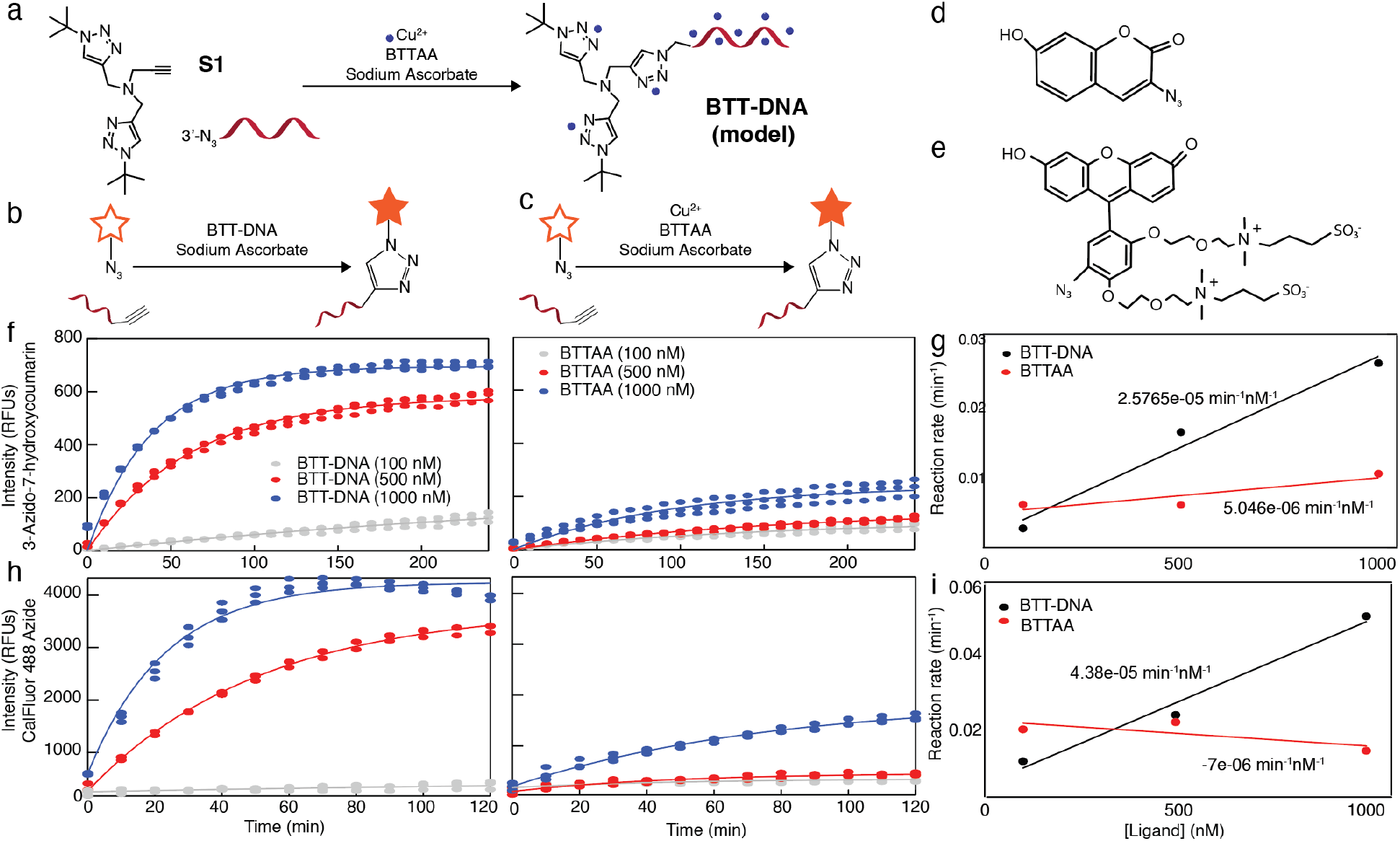
Reaction kinetics of CuAAC enhanced by the BTT-DNA ligand with two different azido fluorogenic dyes. a) Strategy to synthesize BTT-DNA ligand via CuAAC using (S1). b) Schematic showing an alkyne labeled DNA oligonucleotide is ligated to a fluorogenic dye in the presence of Cu^2+^, BTTAA, and sodium ascorbate. c) Schematic showing an alkyne labeled DNA oligonucleotide is ligated to a fluorogenic dye in the presence of BTT-DNA ligand and sodium ascorbate. d) Structure of 3-Azido-7-hydroxycoumarin. e) Structure of CalFluor 488 azide. f) Reaction kinetics of CuAAC using 3-Azido-7-hydroxycoumarin. g) Reaction rate of CuAAC using 3-Azido-7-hydroxycoumarin vs ligand concentration. h) Reaction kinetics of CuAAC using CalFluor 488 azide. i) Reaction rate of CuAAC using CalFluor 488 azide vs. ligand concentration. Each condition has three biological replicates.

### Reaction kinetics of BTT-DNA ligand

The reactivity of the CuAAC reaction in the presence of the new BTT-DNA ligand was determined using a fluorogenic plate-reader assay. We reacted a 15-mer DNA oligomer bearing a 5’ alkyne moiety (henceforth referred to as 5’ alkyne DNA) to a series of azido fluorogenic dyes^28^ (3-Azido-7-hydroxycoumarin and CalFluor 488 azide) using CuAAC in the presence of BTT-DNA and BTTAA (**Fig. 1b,c**). This DNA oligomer sequence is not complementary to any other reaction components. Fluorescence was measured over 2 and 4 hours (**Fig. 1d-i**). BTT-DNA consistently increased activity in accelerating the CuAAC reaction compared to BTTAA for 3-Azido-7-hydroxycoumarin in the nM range. Interestingly, the 100 nM BTT-DNA ligand outperformed the commercial ligand 1.39-fold, demonstrating that BTT-DNA is an effective ligand in the nanomolar range and the absence of exogenous Cu(I). The difference was even more striking at 1000 nM BTT-DNA, at which the fluorescence produced was 3.00-fold higher than the commercial ligand at 2 hours. At 500 nM ligand concentration, we observed that the BTT-DNA ligand outperformed the commercial ligand 4.80-fold (**Fig. 1f**). The rate of CuAAC reaction that was assisted by BTT-DNA ligand is 2.58e^-5^ min^-1^nM^-1^, demonstrating that the CuAAC reaction accelerated by BTT-DNA is significantly faster than the reaction accelerated by the commercial BTTAA ligand with a reaction rate of 5.05e^-6^ min^-1^nM^-1^(**Fig. 1g**).

Next, we performed the fluorogenic plate-reader assay using another fluorogenic dye, CalFluor 488 azide. Upon click reaction in the presence of BTT-DNA ligand, the fluorescence of the CalFluor 488 azide produced at 1,000 nM BTT-DNA was 2.53-fold higher than BTTAA ligand at 2 hours. The BTT-DNA ligand also outperformed the BTTAA ligand 7.54-fold at the concentration of 500 nM (**Fig. 1h**). The most prominent enhancement of the fluorescence was constantly achieved at 500 nM ligand concentration for both fluorogenic dyes. For the CalFluor 488 azide, the rate of CuAAC reaction that was assisted by BTT-DNA ligand is 4.38e^-5^ min^-1^nM^-1^, which is much higher than that of CuAAC reaction assisted by commercial BTTAA ligand (7.0e^-6^ min^-1^nM^-1^; **Fig. 1i**).

### Stability of BTT-DNA ligand after cold storage

To evaluate the stability of the BTT-DNA ligand, we performed the fluorogenic plate-reader assay using 3-Azido-7-hydroxycoumarin in the presence of the BTT-DNA ligand for 4 hours monthly for ten months (**Supplementary Fig. 2**). Using a concentration of 1,000 nM BTT-DNA ligand, we observed that the reaction kinetics were unchanged. There was no significant difference between the fluorescence intensity of each sample across time points over ten months, indicating that our BTT-DNA ligand can be stable even after ten months while stored at -20 °C.

### Contribution of DNA sequence and length in ligand design

We evaluated the reaction with a scrambled single-stranded DNA sequence of equal length using the fluorescent plate-reader assay to assess the DNA sequence’s contribution to the BTT-DNA ligand’s reactivity. We found identical activity, suggesting that the effect is not dependent on the sequence composition (**Supplementary Fig. 3**). Thus, the strategy to synthesize our CuAAC accelerating ligand can be applied to make a BTT-DNA ligand with any sequence. We also varied the length of the DNA oligomer from 15-25 nucleotides to assess the DNA length’s contribution to the BTT-DNA ligand’s reactivity and found that more product formed as the DNA oligomer length increased (**Supplementary Fig. 4**), consistent with the notion that a longer oligomer provides more localized copper for catalyzing the reaction.

### DNA and RNA template-driven proximity ligation with BTT-DNA ligand

We designed a DNA splint that is complementary to both the 5’ alkyne DNA and 3’ BTT-DNA ligand (**Fig. 2a**) to achieve a proximity-mediated click reaction. This design brings the alkyne close to the Cu(I) complexed with the accelerating ligand. We supplemented the reaction with spermine, a polyamine bearing multiple amino groups, and NaCl to improve hybridization (**Supplementary Fig. 5**). To assess the appropriate distance to position the 5’ alkyne on the DNA probe for maximum ligation to the fluorogenic azide, we performed the fluorogenic plate-reader assay for eight different 5’ alkyne DNA probes. For the reaction performed using 3-Azido-7-hydroxycoumarin, the fluorescence intensity produced from the splint in the presence of a DNA alkyne probe with a one nucleotide distance from the BTT-DNA probe was 150.19 RFUs. For the same reaction performed using CalFluor 488 azide, the fluorescence intensity produced was 372.586 RFUs, 1.66-fold higher than the no-splint reaction. This linker distance for the DNA alkyne probe achieved the highest fluorescence enhancement and was used for all proximity experiments (**Supplementary Fig. 5**).

**Fig 2.**
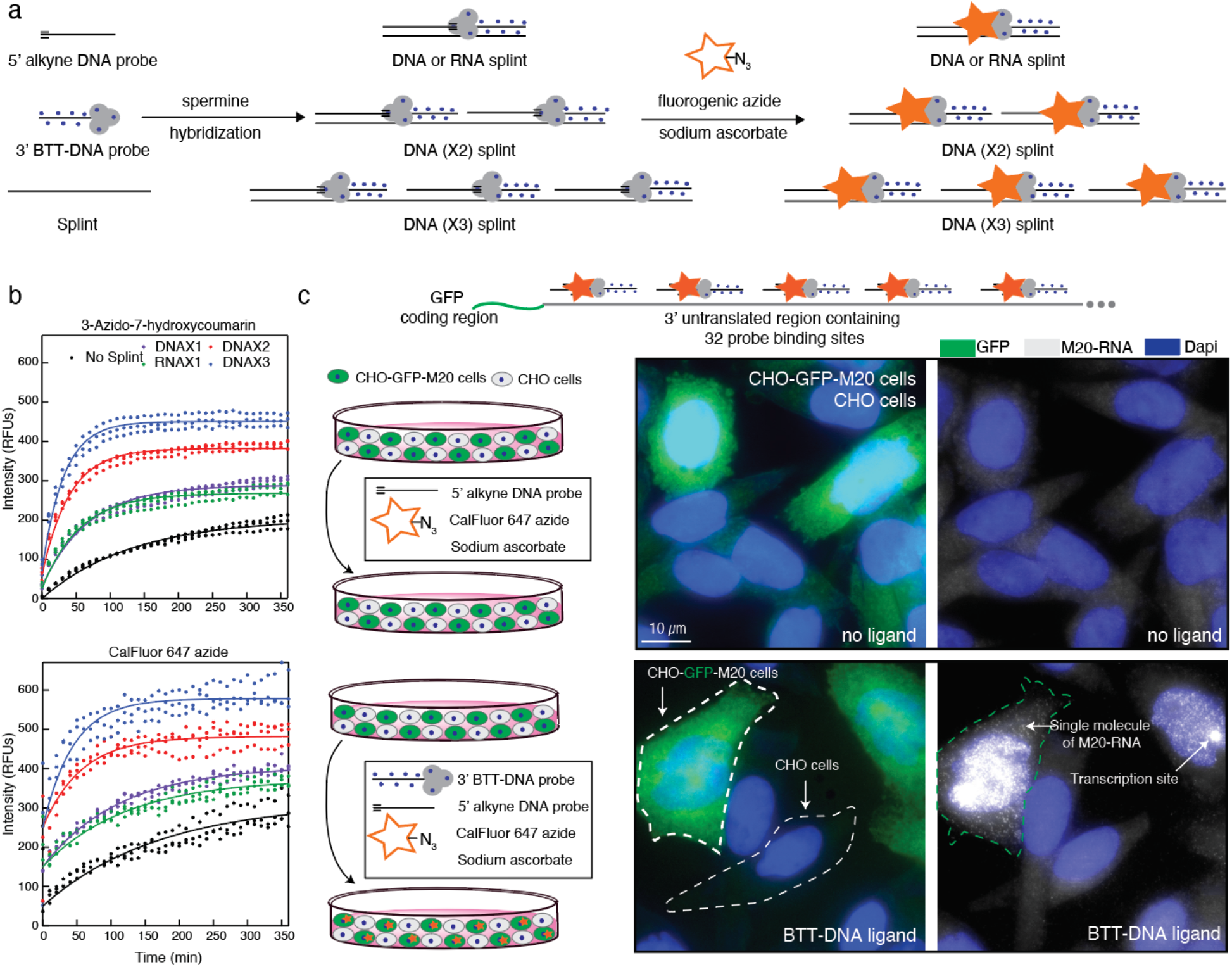
DNA and RNA template-driven proximity ligation with BTT-DNA ligand. a) Schematic of nucleic acid template-driven proximity ligation. b) Reaction kinetics of nucleic acid template-driven proximity ligation with DNA splints harboring different repeats of the binding sequence using 3-Azido-7-hydroxycoumarin and CalFluor 647 azide. Each condition has three biological replicates. c) Microscopy representative of individual RNA molecule detection with nucleic acid template-driven proximity ligation using BTT-DNA. The schematic diagram depicts the proximity click FISH of M20 transgene RNA in transgenic CHO-GFP-M20 cell line. Fixed cells are first treated with 0.2 μM alkyne-modified single-stranded DNA oligonucleotide probe (M20-left-alkyne_20_2), then treated with CuAAC in the presence of CalFluor 647 azide (10 μM), BTT-DNA ligand (20 μM), and sodium ascorbate (2.5 mM) for 1 hour. (white) individual M20 RNA molecule, (green) GFP, (blue) DAPI staining of nuclei.

To enhance the reaction kinetics, we designed DNA splints harboring two and three repeats of the binding sequence (**Fig. 2a**). We assessed the fluorescence after a 6-hour incubation using the fluorogenic plate-reader assay for each condition. For 3-Azido-7-hydroxycoumarin, we found that the total fluorescence produced from a single DNA splint-enhanced reaction is 301.94 RFUs, which is 1.53-fold higher than the fluorescence produced from random collisions (i.e., no splint). As expected, we also found no measurable fluorescence when substituting the targeted BTT-DNA strand with a scrambled BTT-DNA sequence that is not complementary to the DNA splint (**Supplementary Fig. 6**). We also observed that splints harboring more binding sites had higher fluorescence intensities. The fluorescence increased from 301.94 RFUs to 393.76 RFUs and 455.10 RFUs when the binding sites doubled and tripled, respectively (**Fig. 2b**). For the CalFluor 647 azide dye, the total fluorescence produced increased proportionally when we used the DNA splints harboring 1, 2 and 3 repeats of the binding sequence, respectively (**Fig. 2b**). We also found that the RNA splint enhanced reaction kinetics similarly to the DNA splint. For 3-Azido-7-hydroxycoumarin, the fluorescence produced from a single RNA splint-enhanced reaction is 282.85 RFUs, 1.44-fold higher than the fluorescence produced from random collisions. For the CalFuor 647 azide, the total fluorescence increased from 290.77 RFUs to 364.64 RFUs (**Fig. 2b**).

### *In situ* proximity ligation to detect individual mRNAs in fixed cells: no wash smFISH

To test the proximity ligation in fixed cells, we used the transgenic cell line CHO-GFP-M20, which contains a transgene that includes a GFP coding region and 32 tandem copies of the probe-binding sequence (M20) (**Fig. 2c**)^29^. We used the parental CHO cell line which does not contain the transgene as a negative control. First, we optimized the CuAAC reaction time to bind an azido dye to 5’ alkyne DNA probe in fixed CHO-GFP-M20 cells. We optimized the CuAAC reaction time and achieved the best labeling by performing a 1-hour CuAAC reaction (**Supplementary Fig. 7**).

After CuAAC reaction time optimization, we performed the CuAAC reaction using the BTT-DNA ligand with eight different 5’ alkyne DNA probes for targeting the M20 transgene sequence to find the best 5’ alkyne DNA probe for maximum ligation to the fluorogenic azide for transgene detection in CHO-GFP-M20 cells (**Supplementary Fig. 8**). Clean and robust labeling of transcription sites in the nucleus and individual RNA molecules in the nucleus and cytoplasm was observed via CuAAC in the presence of a 5’ alkyne DNA probe with a two-nucleotide distance from the BTT-DNA probe (**Fig. 2c, Supplementary Fig. 8**). Then, we cultivated an equal mix of CHO-GFP-M20 and parental CHO cells together. The cells were fixed and permeabilized, then reacted with CalFluor 647 azide via CuAAC in the presence of BTT-DNA ligand for 1 hour. We observed specific labeling of transcription sites in the nucleus and individual RNA molecules in the nucleus and cytoplasm (**Fig. 2c**). The cells with bright puncta in the CalFluor 647 azide channel colocalize with GFP-positive cells, indicating that the probe binds specifically (**Fig. 2c**). In short, we achieved the detection of individual mRNAs specifically using a fluorogenic dye and non-exogenous copper required proximity click RNA FISH approach assisted by the BTT-DNA ligand.

### Fixed cell detection of biomolecules with BTT-DNA-assisted CuAAC

Next, we tested the activity of our BTT-DNA ligand in a cellular environment to detect various biomolecules in fixed cells. 5-Ethynyl-2’-deoxyuridine (EdU) is an alkyne-derivatized thymidine analog that may be metabolically incorporated into newly synthesized DNAs during active DNA synthesis in live cells^4^. Following overnight metabolic incorporation, cells were fixed and permeabilized, then reacted with CalFluor 647 azide via CuAAC. The negative control containing no ligand showed very low background fluorescence. We compared the performance of the BTT-DNA ligand to commercially available BTTAA (**Supplementary Fig. 9**). We observed robust labeling of nascent DNA in the nucleus using the CuAAC accelerated by our BTT-DNA ligand (**Fig. 3a, Supplementary Fig. 10;** 3.12-fold enhancement of signal over background), while the fluorescence of the cell nucleus using the CuAAC assisted by CuSO_4_ and BTTAA was very low.

**Fig 3.**
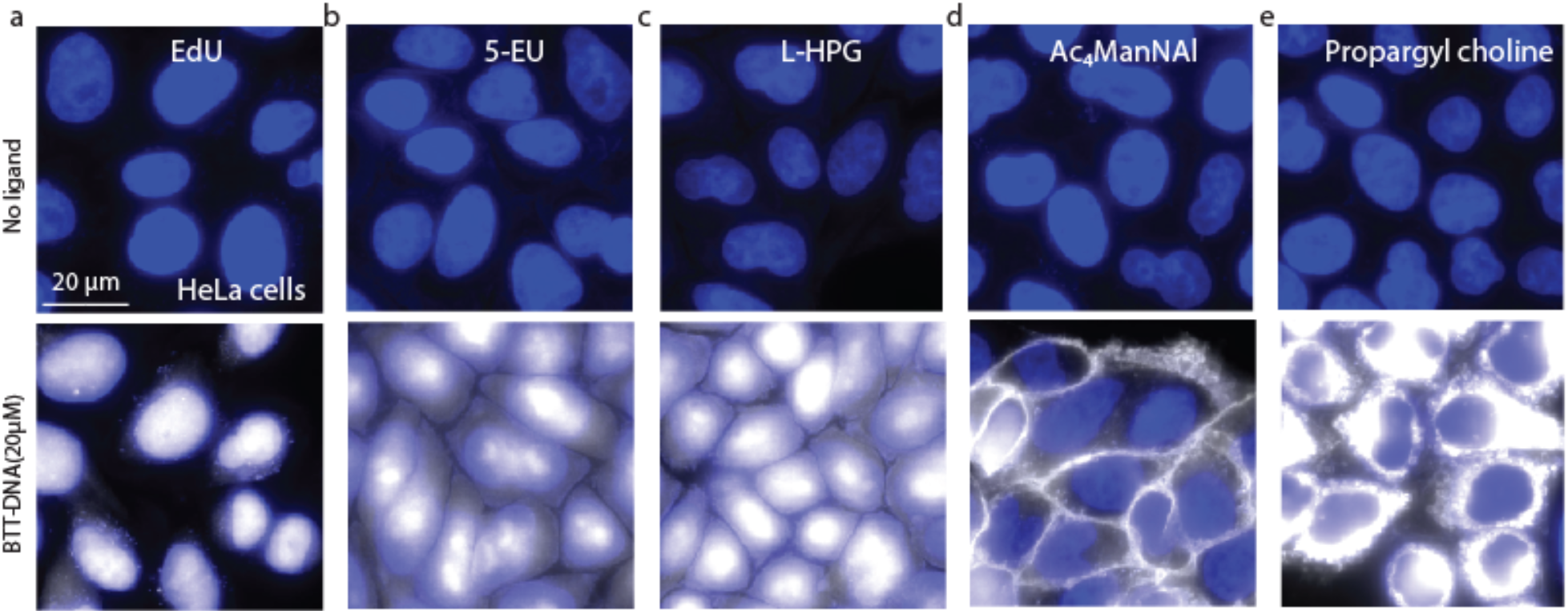
Detection and labeling of the intracellular and extracellular biomolecules using CuAAC in the presence of BTT-DNA on fixed cells. a) Representative fluorescence microscopy for detection of nascent EdU-labeled DNA via CuAAC in the presence of BTT-DNA ligand on fixed cells. b) Representative fluorescence microscopy (two biological replicates) of detection of nascent 5-EU-labeled RNA via CuAAC in the presence of BTT-DNA ligand on fixed cells. c) Representative fluorescence microscopy (two biological replicates) of detection of nascent L-HPG-labeled protein via CuAAC in the presence of BTT-DNA ligand on fixed cells.d) Representative fluorescence microscopy (two biological replicates) of detection of nascent Ac_**4**_MaNAl-labeled sialic acid via CuAAC in the presence of BTT-DNA ligand on fixed cells. e) Representative fluorescence microscopy (two biological replicates) of detection of nascent propargyl choline-labeled phospholipids via CuAAC in the presence of BTT-DNA ligand on fixed cells. On the top row, fixed cells are treated with CuAAC in the presence of 10 μM CalFluor 647 azide and 2.5 mM sodium ascorbate; on the bottom row, fixed cells are treated with CuAAC in the presence of 10 μM CalFluor 647 azide, 20 uM BTT-DNA ligand, and 2.5 mM sodium ascorbate. (white) CalFluor 647 azide labeling of nascent intracellular and extracellular biomolecules, (blue) DAPI staining of nuclei.

5-Ethynyl Uridine (EU) is an alkyne-derivatized uridine analog that may be metabolically incorporated into nascent RNAs in live cells^19^. Following a three-hour metabolic incorporation, cells were fixed and permeabilized, then reacted with CalFluor 647 Azide via CuAAC, comparing the performance of BTT-DNA ligand to commercially available BTTAA (**Fig. 3b, Supplementary Fig. 9**). For the cells treated with 20 μM BTT-DNA ligand, we observed strong fluorescence intensity in the cell nuclei, especially in the nucleoli ^19,30^(**Fig. 3b**). The total fluorescent signal produced in the nucleus in the presence of the BTT-DNA ligand was 2.18-fold higher than that in the no ligand control (**Fig. 3b, Supplementary Fig. 10**). In contrast, cells treated with CuSO_4_ and BTTAA had minimal fluorescence signals.

L-homopropargylglycine (L-HPG) is a cell-permeable alkyne probe that can be metabolically incorporated into nascent proteins in methionine-starved cells^28^. We treated cells for 2 hours with L-HPG, then fixed and permeabilized. We then reacted these samples with CalFluor 647 Azide via CuAAC in the presence of BTT-DNA to visualize the newly synthesized proteins (**Fig. 3c, Supplementary Fig. 9,10**). We found strong fluorescence signals in the cell nucleus using the CuAAC assisted by 20 μM BTT-DNA ligand, which shows a 2.60-fold enhancement of fluorescence compared to the ligand-less ones (**Fig. 3c, Supplementary Fig. 10**).

Further, we metabolically labeled HeLa cells with Ac_**4**_ManNAl to introduce an alkyne tag to cell surface sialic acids^31^, clicked a fluorophore to the alkynyl sugar, and imaged using fluorescence microscopy (**Fig. 3d, Supplementary Fig. 9,10**). For the cells treated with 20 μM BTT-DNA ligand, a strong fluorescent signal was observed on the cell membrane where the metabolically labeling sialylated glycans are located (**Fig. 3d**). Compared to the control, cells treated with 20 μM BTT-DNA showed 9.09-fold enhancement of signals (**Fig. 3d, Supplementary Fig. 10**).

Next, we metabolically incorporated propargyl-choline into newly synthesized phospholipids. Cells were incubated with propargyl-choline for 24 hours, then fixed and permeabilized, and then stained with CalFluor 647 Azide via CuAAC in the presence of BTT-DNA to visualize the newly synthesized phospholipids (**Fig. 3e, Supplementary Fig. 9,10**). The cells showed intense staining in the cellular membrane and the intracellular structure ^30^ (**Fig. 3e**). Remarkably, the total fluorescence produced in the cytoplasm in the presence of BTT-DNA was 239.87-fold higher than that in the no ligand control (**Fig. 3e, Supplementary Fig. 10**).

### BTT-DNA ligand enables the detection of biomolecules on the surface of live cells

To test the activity of BTT-DNA in driving the CuAAC reaction for live cells, we metabolically labeled HeLa cells with Ac_**4**_ManNAl (**Fig. 4a, Supplementary Fig. 11**), then clicked CalFluor 647 azide to the alkynyl sugar on the cell surface while the cells were incubated on ice to prevent internalization of the reagents. The BTT-DNA accelerated CuAAC produced a robust fluorescent signal on the cell membrane, showing a 7.26-fold enhancement of fluorescent signal over the no ligand background (**Fig. 4a, b**).

**Fig 4.**
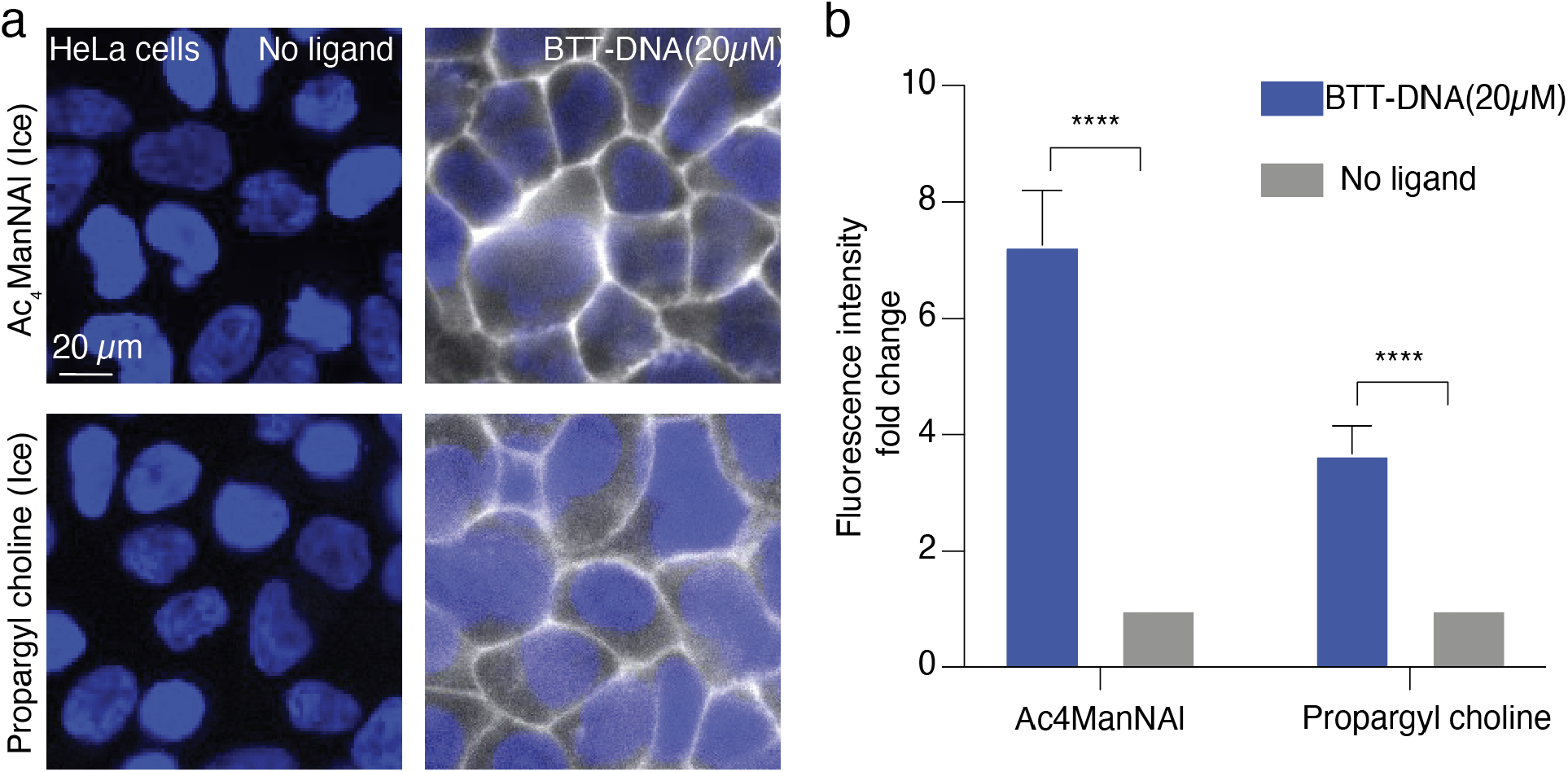
Representative fluorescence microscopy (n=2) of the cell-surface biomolecule labeling and detection in the presence of BTT-DNA on live cells. a) Metabolic labeling and detection of cell-surface sialic acids and choline-containing phospholipids with BTT-DNA on live cells. On the left column, live cells are treated with CuAAC in the presence of 10 μM CalFluor 647 azide and 2.5 mM sodium ascorbate; on the right column, live cells are treated with CuAAC in the presence of 10 μM CalFluor 647 azide, 20 uM BTT-DNA ligand, and 2.5 mM sodium ascorbate. (white) CalFluor 647 azide labeling of sialic acids and choline-containing phospholipids, (blue) Hoechst 33342 nuclei staining. b) Quantifying normalized signal over background for the fluorescence microscopy assisted by BTT-DNA in panel (a).

Choline-containing phospholipids were also detected by metabolic incorporation of the propargyl-choline to evaluate the activity of our BTT-DNA ligand for extracellular labeling in live cells (**Fig. 4a, Supplementary Fig. 11**). We found robust labeling of the newly synthesized phospholipids in the cell membrane (**Fig. 4a, b**), which is 3.67-fold higher than that of the no-ligand control.

### Characterizing copper-induced cellular toxicity in the presence of BTT-DNA ligand

To evaluate the cytotoxicity of copper with the new Cu(I) accelerating ligand BTT-DNA, we compared HeLa cells that were treated with CuSO_4_ at the copper concentration (300 μM) to HeLa cells treated with BTT-DNA ligand in the equivalent copper ion concentration (30 μM) (each molecule of ligand is complexed to ∼10 copper ions; **Fig. 1**). Each of these was delivered under reducing conditions with sodium ascorbate, whose optimal concentration (2 mM; **Supplementary Fig. 12**) and treatment time (5 hours; **Supplementary Fig. 13**) were necessary to minimize cell toxicity. We also optimized the concentration of lipofectamine RNAiMAX transfection reagents to eliminate the fluence on cell biology (0.5 μl; **Supplementary Fig. 14**). We observed that most cells treated with free CuSO_4,_ and sodium ascorbate were “blebbing” (i.e., cell membrane bulges out indicative of apoptosis). In contrast, the morphology of cells treated with only sodium ascorbate, or a combination of BTT-DNA and sodium ascorbate were comparable to the negative control (**Fig. 5a**). This suggests that BTT-DNA ligand can prevent cellular “blebbing” induced by Cu(I).

**Fig 5.**
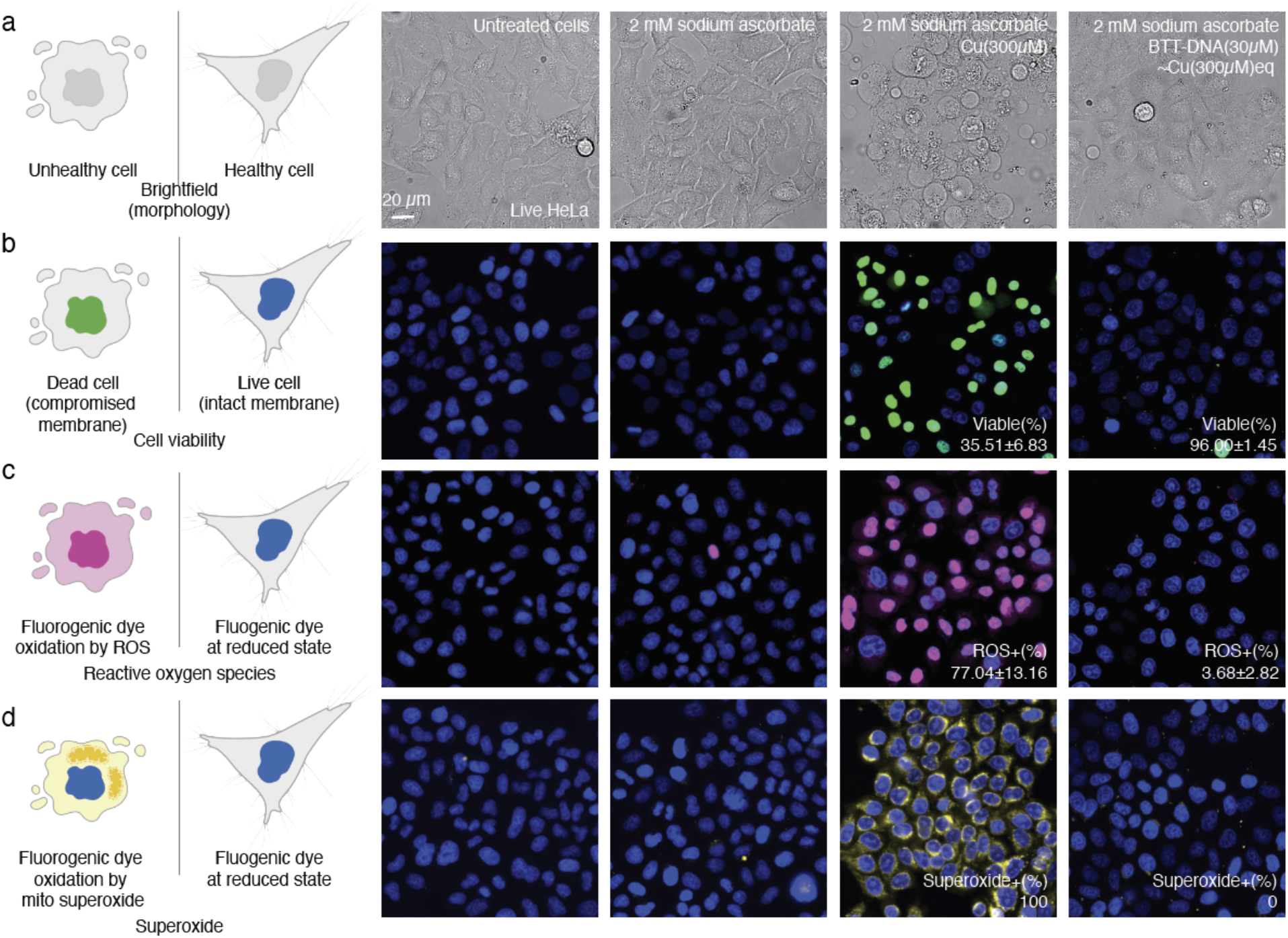
Biocompatibility of the BTT-DNA ligand (n=2). a) Representative micrographs of cell morphology comparing BTT-DNA ligand to copper on Hela cells (scale bar applies to all images). b) Representative micrographs of cell viability assay comparing BTT-DNA ligand to copper on Hela cells. c) Representative micrographs of cell reactive oxygen species comparing BTT-DNA ligand to copper on Hela cells. d) Representative micrographs of cell superoxide comparing BTT-DNA ligand to copper on Hela cells. (blue) Hoechst 33342 staining of nuclei of all cells, (green) SYTOX green nucleic acid staining of nuclei of dead cells. (red) CellROX Green Reagent of reactive oxygen species of nuclei of all cells, (yellow) MitoSOX™ green indicators of mitochondrial superoxide of all cells.

Next, we compared the viability of the treated cells using the ReadyProbes cell viability imaging kit. The nuclei of cells with compromised membranes are stained with the SYTOX green and measured with fluorescence microscopy (**Fig. 5b, Supplementary Fig. 15**). We calculated the viability for cells treated with Cu(I) to be 35.51±6.83% viable. In contrast, the viability ratio for cells treated with BTT-DNA ligand is 96.00 ±1.45% (**Fig. 5b**). This strengthens the claim that the BTT-DNA ligand protects against Cu(I) induced toxicity.

A major concern for applying CuAAC into biological systems is the formation of reactive oxygen species (ROS), which can cause oxidative damage to proteins ^32^. To compare ROS generated from the treatment of live cells with BTT-DNA ligand, we measured the amount of the ROS in live cells by CellROX Green Reagent, which fluoresces after oxidation by ROS (**Fig. 5c, Supplementary Fig. 15**). We found that 77.04 ± 13.16% of cells were positive for ROS staining after free Cu(I) treatment. In contrast, the ratio for cells treated with BTT-DNA was only 3.68 ± 2.82% (**Fig. 5c**), indicating that BTT-DNA significantly reduces the ROS produced by Cu(I) delivery in live cells. We also used the MitoSOX green superoxide indicator. This fluorogenic dye can readily oxidize by superoxide produced by mitochondria rather than ROS (**Fig. 5d, Supplementary Fig. 15**). For cells treated by Cu(I), all cells showed fluorescence in the cytoplasm, indicating superoxide production. All cells treated by BTT-DNA were negative for superoxide (**Fig. 5d**). Together, these results indicate that BTT-DNA has a protective effect from Cu(I) toxicity and prevents the formation of ROS and superoxide.

### BTT-DNA ligand enables the detection of intracellular phospholipids on live cells

After a demonstration of the biocompatibility of our BTT-DNA ligand, we performed the intracellular biomolecule labeling via CuAAC assisted by our BTT-DNA ligand. We started with the metabolic incorporation of propargyl-choline into newly synthesized phospholipids by 24 hours of propargyl-choline treatment, through which the endoplasmic reticulum (ER), the Golgi, the mitochondrial and plasma membranes were labeled with choline-containing phospholipids^30^. To ensure that our BTT-DNA ligand enters the cells to accelerate the CuAAC reaction, we used lipofectamine RNAiMAX transfection reagent, which has high transfection efficiency for siRNA. Since lipofectamine RNAiMAX encapsulates double-stranded nucleic acid, and our BTT-DNA is single-stranded, we designed a short single-stranded DNA that is complementary to the sequence of BTT-DNA and hybridized to form a duplex before transfection.

After encapsulating the BTT-DNA duplex with the liposome, 80 μM CalFluor 647 azide was added to the mixture. This concentration of dye leads to detectable passive transport of dye into the cytoplasm while maintaining a sufficiently low dye background in live cells (**Supplementary Fig. 16**). We also optimized the transfection time to 4 hours to transport the duplex into the cytoplasm of cells (**Supplementary Fig. 17**). After 4 hours of transfection, we washed the cells to ensure that CuAAC occurs exclusively inside the cells and not on extracellular alkynes, followed by a 30-minute sodium ascorbate incubation (**Fig. 6a**). We observed strong fluorescence intensity in the cytoplasm and membrane of cells treated with BTT-DNA (**Fig. 6b**). We also conducted time-lapse imaging of intracellular labeling of phospholipids for 2 hours. We observed that some cells that were non-fluorescent at 30 minutes began to exhibit fluorescence at 1 hour and became more fluorescent at 2 hours (**Fig. 6c**). We also showed that the fluorescence signal increased significantly over time, and the fluorescence enhancement compared to that at 30 minutes is 1.454±0.159 fold, 1.738±0.231 fold and 1.973±0.277 fold at 60 minutes, 90 minutes and 120 minutes, respectively (**Fig. 6c, d**). Cell viability measurements indicated that almost 100% of the cells were viable (**Supplementary Fig. 18**), indicating that the BTT-DNA ligand enables intracellular labeling of phospholipids in live cells. Additionally, to confirm the presence of free choline-containing phospholipids on the cell surface, we conducted a CuAAC reaction using Alexa Fluor 555 azide, which is a fluorescent dye with BTTAA-CuSO_4_ complex for 5 minutes after intracellular phospholipids labeling. We observed colocalization of AF555 with CF647 (**Fig. 6e**), demonstrating the presence of the choline-containing phospholipids on the cell surface and consistent with the model in **Figure 6a**.

**Fig 6.**
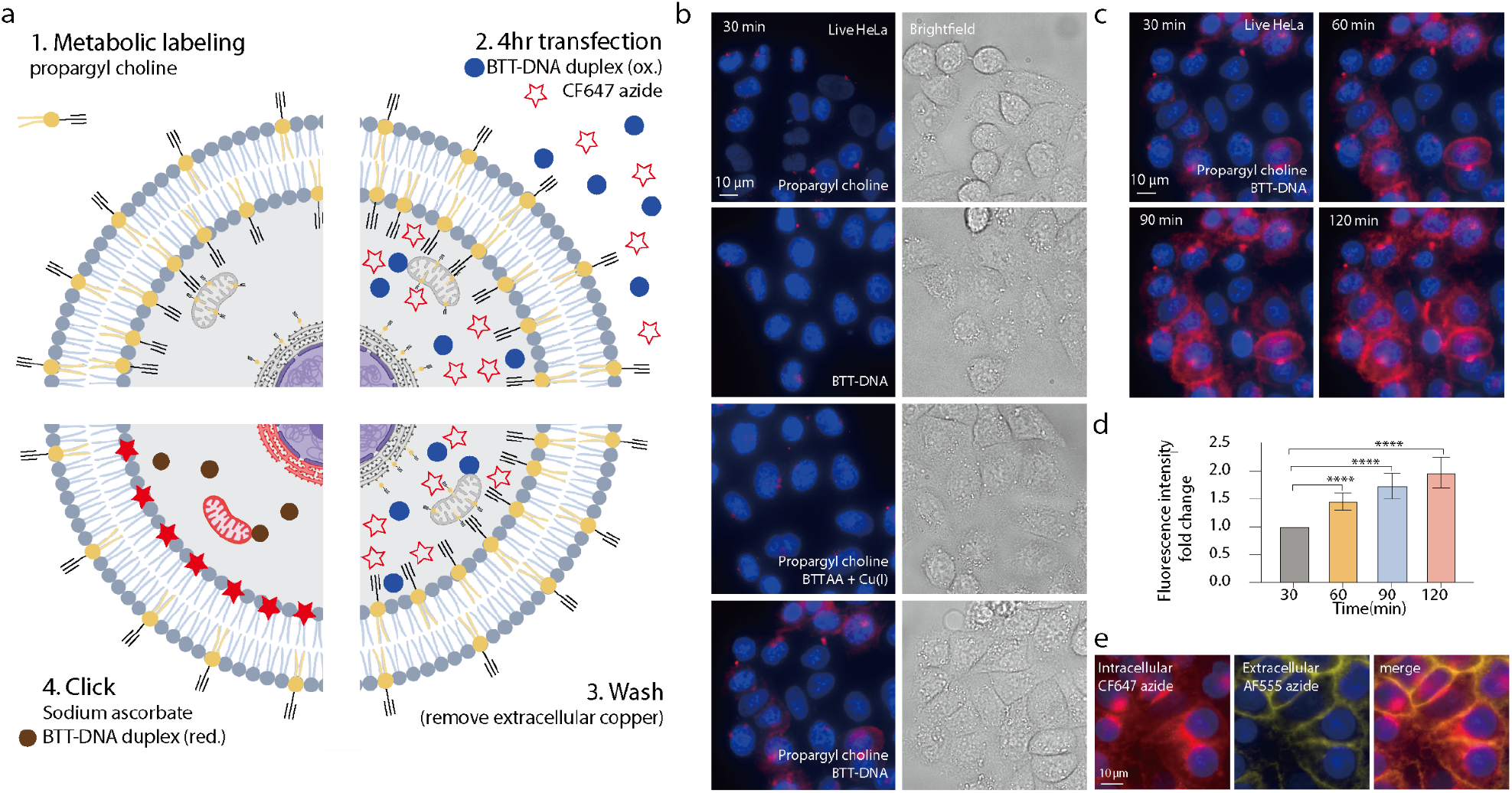
Intracellular phospholipid labeling and detection in the presence of BTT-DNA on live cells. a) Workflow of metabolic labeling and detecting intracellular choline-containing phospholipid with BTT-DNA on live cells. (ox.) BTT-DNA duplex with oxidized copper (red.) BTT-DNA duplex with reduced copper. b) Representative fluorescence micrographs (n=3) of the intracellular phospholipids labeling and detection in the presence of BTT-DNA on live cells. (from top to bottom) Live cells are treated with CuAAC in the presence of 1) propargyl choline, lipofectamine RNAiMAX transfection reagent, CalFluor 647 azide and sodium ascorbate, 2) lipofectamine RNAiMAX transfection reagent, BTT-DNA ligand, CalFluor 647 azide and sodium ascorbate, 3) propargyl choline, lipofectamine RNAiMAX transfection reagent, BTTAA, CuSO4, CalFluor 647 azide and sodium ascorbate, 4) propargyl choline, lipofectamine RNAiMAX transfection reagent, BTT-DNA ligand, CalFluor 647 azide and sodium ascorbate. (red) CalFluor 647 azide labeling of choline-containing phospholipid, (blue) Hoechst 33342 nuclei staining. c) Time-lapse fluorescence microscopy of the intracellular phospholipids labeling in BTT-DNA presence for 2 hours. Live cells are treated with CuAAC in the presence of propargyl choline, lipofectamine RNAiMAX transfection reagent, BTT-DNA ligand, CalFluor 647 azide, and sodium ascorbate. d) Fluorescence intensity fold-change enhancement as a function of reaction time for the BTT-DNA experiments in panel (c). Error bars represent the standard deviation of 60 cells from three replicates. * p≤ 0.05, ** p≤ 0.01, *** p≤ 0.001, and **** p≤ 0.0001. T-test was used. e) Representative fluorescence microscopy of the intracellular phospholipids labeling and detection in the presence of BTT-DNA and extracellular phospholipids labeling and detection in the presence of BTTAA-CuSO_4_ complex on live cells. (red) CalFluor 647 azide labeling of choline-containing phospholipid, (yellow) Alexa Fluor 555 azide labeling of choline-containing phospholipid on the surface, (blue) Hoechst 33342 nuclei staining.

## Discussion

Here, we developed BTT-DNA, a DNA-conjugated CuAAC accelerating ligand that reduces the need to add free copper for CuAAC-mediated intracellular labeling while maintaining fast reaction kinetics. Importantly, this ligand eliminates the need to add copper salt to the CuAAC reaction because it is already complexed with the ligand. With this new CuAAC accelerating ligand, the total ligand concentration necessary for the CuAAC *in vitro* may be decreased to the nM range while remaining complexed to an average of 10 copper ions per ligand with different fluorogenic azido dyes. This new ligand also significantly drove forward the CuAAC reaction, showing more robust intracellular labeling in fixed cells and achieving live cell detection of biomolecules in the cell surface and cytoplasm.

Previous studies have shown DNA splint-enhanced CuAAC^33–35^ by combining the azide and the alkyne probes; however, these systems preclude using fluorogenic dyes, which are ideal for live-cell applications. The BTT-DNA ligand may further drive the CuAAC reaction kinetics when hybridized to a DNA or RNA splint close to an alkyne-modified probe and clicked to a fluorogenic azide. This demonstrated that a completely non-fluorescent system may be used to drive the CuAAC reaction kinetics of a fluorogenic dye, which is highly desirable for live-cell imaging. The reaction was enhanced by adding spermine and NaCl before CuAAC. We suspect that the spermine contributes to the DNA-DNA duplex and DNA-RNA duplex stability, which is vital for the subsequent CuAAC reaction. Additionally, spermine can help mask the negative charge of oligonucleotides^37^, which may sequester the copper away from the ligation site. We also showed that DNA splints with multiple binding sites could amplify the total fluorescence produced. Our BTT-DNA ligand could also enhance the RNA template-driven proximity ligation *in situ* on fixed cells.

Our experiments demonstrated that the BTT-DNA ligand could be used for standard CuAAC ligation assays in cellular environments to detect extracellular and intracellular biomolecules in fixed cells. For the detection of nascent DNAs in the cell nucleus using EdU labeling, we observed that the cells treated with 20 μM BTT-DNA ligand showed much stronger staining not only than cells treated without ligand but also than cells treated with 20 μM BTTAA commercial ligand, demonstrating that BTT-DNA ligand outperformed the commercial reagent^9^ in detection of nascent DNAs in fixed cells. Robust labeling of nascent RNAs in the cell nuclei in the presence of BTT-DNA ligand was observed compared to the same concentration of commercial BTTAA ligand. As for the protein detection using L-HPG labeling, we found bright fluorescent signals in the nucleus and cytoplasm in the presence of BTT-DNA. For all three intracellular biomolecules, there was no significant difference in the fluorescence signal to the background using CuAAC accelerated by BTT-DNA compared to the no ligand control, indicating that the BTT-DNA ligand can consistently outperform the commercial ligand in biological environments with lower concentration. Interestingly, we achieved the highest signal-to-background ratio in phospholipid detection using propargyl choline labeling. This demonstrates that the BTT-DNA ligand can outperform the BTTAA ligand with a high yield.

We demonstrated that the BTT-DNA ligand significantly lowered the copper-induced cytotoxicity. When the cells were treated with free copper and sodium ascorbate for 5 hours, the morphology of cells changed, and almost all cells were “blebbing,” indicative of pre-apoptosis. We showed that copper in the presence of sodium ascorbate could compromise the cell membrane integrity and cause cell death. Interestingly, we also observed compromised cell membranes with sodium ascorbate alone when we increased the treatment time to 6 hours. We found that not every blebbing or round cell showed fluorescence for SYTOX green, and cells with normal morphology could also show fluorescence for SYTOX green, which means blebbing is not the only indicator of the compromised cell membrane.

Copper-treated cells also produce oxidative stress, which can damage the biological material in cells, like ROS, which can oxidize the CellRox green reagent. This reagent is a fluorogenic probe that is non-fluorescent in the reduced state and exhibits bright green photostable fluorescence after being oxidized by the ROS and bound to DNA. We saw extremely strong fluorescence in the nucleus while fluorescence was dimmer in the cytoplasm. We also saw strong fluorescence in the nucleus of cells treated with 2.5 mM sodium ascorbate, indicating that a high concentration of sodium ascorbate could also cause the production of ROS in live cells. Superoxide is a form of oxidative stress responsible for damaging biomolecule structure integrity. MitoSOX green superoxide indicator measures the amount of the superoxide produced only by mitochondria. We saw strong fluorescence in the cytoplasm for cells treated with copper in the presence of sodium ascorbate. The cells treated with BTT-DNA ligand appeared morphologically like the negative control for all tests. This indicates that our BTT-DNA ligand is biocompatible with living cells by lowering copper-induced cytotoxicity.

Finally, we demonstrated that the BTT-DNA ligand may be used for intracellular detection of propargyl choline-labeled nascent phospholipids on live cells. This allowed us to track intracellular choline-containing phospholipids via CuAAC assisted by our BTT-DNA ligand for 2 hours. Our live cell labeling scheme could enable future applications such as tracking the dynamics of other major biomolecules like lipid metabolism in response to drug treatment or tracking of global cellular transcription over time. Overall, the newly developed CuAAC accelerating ligand allows sensitive detection of biomolecules in fixed and live cells and holds great promise for further application of CuAAC for intracellular, live-cell detection.

## Materials and Methods

### Synthesis and purification of BTT-DNA ligand

A 120-fold molar excess of *N,N*-bis((1-*tert*-butyl-1*H*-1,2,3-triazol-4-yl)methyl)prop-2-yn-1-amine (S1; purchased from the AECOM chemical biology core facility) was reacted with 3’ N_3_-labeled 15 mer ssDNA oligo using BTTAA ligand-assisted CuAAC ([BTTAA]:[CuSO_4_]=2:1) (See **Supplementary methods)**. The reaction mixture was shaken at 600 rpm for 30 mins at 37°C, resulting in a crude BTT-DNA ligand. The BTT-DNA ligand was purified using 3.5kD MWCO dialysis tubing for 24 hours in 3.5 L of water at room temperature (Repligen). Invitrogen Qubit ssDNA Assay Kit was used to measure the concentration of the purified BTT-DNA ligand.

### Inductively Coupled Plasma Mass Spectrometry (ICP-MS)

To assess the copper complexed with the BTT-DNA ligand, we prepared the ligand for ICP-MS. The sample was dissolved (0.1 ml) in 0.9 ml of concentrated HNO_3_. The sample was heated for an hour at 90°C. Then, the temperature was increased to 150°C, and the sample was boiled. 0.5 ml of concentrated HNO3 was added to the sample twice during the boiling period. When the sample was completely dried, we added 0.5 ml of concentrated HNO_3_ and diluted it with NF-H_2_O to bring the final volume to 10 ml. To confirm the accuracy of our ICP-MS data, we submitted two replicates. Samples were submitted to Robertson Microlit Laboratories at Ledgewood, New Jersey, for ICP-MS.

### Tissue culture conditions

HeLa and CHO cells were grown in Dulbecco’s modified Eagle’s medium (DMEM) supplemented with 10% FBS. All cells were incubated in a 5% CO2, water-saturated incubator at 37°C.

### Proximity click FISH of M20 transgene RNA on fixed cells

Cho cells and CHO-GFP-M20 cells were cultured in Dulbecco’s modified Eagle’s medium (DMEM) supplemented with 10% FBS for three days and harvested by centrifugation (300Xg,3 min) and resuspended in fresh media and seeded into the 8-well chamber (Fisher scientific). We seeded an equal concentration of Cho cells and CHO-GFP-M20 cells together, and the total cell number is 60000 cells/well, and the total volume per well is 300 μl and incubated for 24 hours.

Cells were rinsed with 1X PBS and fixed with 4% formaldehyde for 10 minutes, then permeabilized with 70% EtOH in NF-H_2_O at 4°C overnight. Cells were rinsed with 1X PBS/10% formamide, followed by treatment with 100 μl smFISH hybridization buffer (10% formamide and 10% dextran sulfate in 1X PBS) containing 2 μl of M20-left-alkyne_20_2(10 μM) for 24 hours at 37°C. After hybridization, cells were washed with 300 μl 1X PBS/10% formamide for 30 minutes at 37°C. Then, it was treated with 10 μM CalFluor 647 Azide, BTT-DNA ligand ([BTT-DNA] = 20μM), and 2.5 mM freshly prepared sodium ascorbate for 1 hour at 37°C. After the click reaction, the cells were rewashed with 300 μl 1X PBS/10% formamide for 30 minutes twice at 37°C and counterstain with DAPI stain diluted with 1X PBS/10% formamide for 20 minutes at 37°C. Then, 1X PBS was added to each well to prepare for imaging. For negative control, we treated the cells with M20-left-alkyne_20_2 without ligand.

### 5-ethynyl-2’-deoxyuridine (EdU) labeling of nascent DNAs on fixed cells

Hela cells were cultured in Dulbecco’s modified Eagle’s medium (DMEM) supplemented with 10% FBS for three days, harvested by centrifugation (300Xg,3 min), resuspended in fresh media, and seeded into the 18-well chamber (Cellvis). 1.5 μl EdU (1mM; Thermo Fisher Scientific) was added to each well, and the total volume per well was 150 μl, and incubated for 24 hours.

Cells were rinsed with 1X PBS and fixed with 4% formaldehyde for 10 minutes, then permeabilized with 70% EtOH in NF-H_2_O at 4°C overnight. Cells were washed with 1X PBS/10% formamide for 30 minutes at 37°C, followed by treatment with 10 μM CalFluor 647 Azide, premixed BTTAA-CuSO_4_ complex ([BTTAA]:[CuSO_4_] = 2:1,[CuSO_4_] = 10 μM) or BTT-DNA ligand ([BTT-DNA] = 20μM) and 2.5 mM freshly prepared sodium ascorbate for 30 minutes at 37°C. After the click reaction, the cells were rewashed with 150 μl wash buffer for 30 minutes twice at 37°C and counterstain with DAPI stain diluted with 1X PBS/10% formamide for 20 minutes at 37°C. Then, 1X PBS was added to each well to prepare for imaging.

### 5-ethynyl uridine (EU) labeling of nascent RNAs on fixed cells

Hela cells were cultured in Dulbecco’s modified Eagle’s medium (DMEM) supplemented with 10% FBS for three days, harvested by centrifugation (300Xg,3 min), resuspended in fresh media, and seeded into the 18-well chamber (Cellvis). The total volume per well is 150 μl. After one day of incubation at 37°C, 1.5 μl 5-ethynyl uridine (EU) was added to each well to the 1mM final concentration and returned to the incubator for three hours of incubation.

Cells were rinsed with 1X PBS and fixed with 4% formaldehyde for 10 minutes, then permeabilized with 70% EtOH in NF-H_2_O at 4°C overnight. Cells were washed with 1X PBS/10% formamide for 30 minutes at 37°C, followed by treatment with 10 μM CalFluor 647 Azide, premixed BTTAA-CuSO_4_ complex ([BTTAA]:[CuSO_4_] = 2:1,[CuSO_4_] = 10 μM) or BTT-DNA ligand ([BTT-DNA] = 20μM) and 2.5 mM freshly prepared sodium ascorbate for 30 minutes at 37°C. After the click reaction, the cells were rewashed with 150 μl wash buffer for 30 minutes twice at 37°C and counterstained with DAPI stain diluted with 1X PBS/10% formamide for 20 minutes at 37°C. Then, 1X PBS was added to each well to prepare for imaging.

### L-homopropargyl(L-HPG) labeling of nascent proteins on fixed cells

Hela cells were cultured in Dulbecco’s modified Eagle’s medium (DMEM) supplemented with 10% FBS for three days, harvested by centrifugation (300Xg,3 min), and resuspended in fresh media and seeded into the 18-well chamber (Cellvis). The total volume per well is 150 μl. After one day of incubation at 37°C, the cell growth medium was removed and replaced with a methionine-free DMEM medium for 30 minutes to eliminate the methionine. Then 1.5 μl L-HPG was added to each well to the 2 mM final concentration and returned to the incubator for 2-hour incubation.

Cells were rinsed with 1X PBS and fixed with 4% formaldehyde for 10 minutes, then permeabilized with 70% EtOH in NF-H_2_O at 4°C overnight. Cells were washed with 1X PBS/10% formamide for 30 minutes at 37°C, followed by treatment with 10 μM CalFluor 647 Azide, premixed BTTAA-CuSO_4_ complex ([BTTAA]:[CuSO_4_] =2:1,[CuSO_4_] = 10 μM) or BTT-DNA ligand ([BTT-DNA] = 20μM) and 2.5 mM freshly prepared sodium ascorbate for 30 minutes at 37°C. After the click reaction, the cells were rewashed with 150 μl wash buffer for 30 minutes twice at 37°C and counterstained with DAPI stain diluted with 1X PBS/10% formamide for 20 minutes at 37°C. Then, 1X PBS was added to each well to prepare for imaging.

### N -(4-pentynoyl) mannosamine (Ac4MaNAl) labeling of cell-surface sialic acid on fixed cells

Hela cells were cultured in Dulbecco’s modified Eagle’s medium (DMEM) supplemented with 10% FBS for three days, then harvested and resuspended in fresh media. 2.5 μl of Ac_4_ManNAl (5mM; Click Chemistry Tools) was added to each well of the 18-well chamber (Cellvis) and dried for 20 min. Cells were seeded into the Ac_4_ManNAl-treated 18-well chamber, and the total volume per well was 150 μl and incubated for 24h.

Cells were rinsed with 1X PBS and fixed with 4% formaldehyde for 10 minutes. Cells then were washed with 1X PBS/10% formamide for 30 minutes at 37°C, followed by treatment with 10 μM CalFluor 647 Azide, premixed BTTAA-CuSO_4_ complex ([BTTAA]:[CuSO_4_] = 2:1,[CuSO_4_]= 10 μM) or BTT-DNA ligand ([BTT-DNA] = 20μM) and 2.5 mM freshly prepared sodium ascorbate for 30 minutes at 37°C. After the click reaction, the cells were rewashed with 150 μl wash buffer for 30 minutes twice at 37°C and counterstain with DAPI stain diluted with 1X PBS/10% formamide for 20 minutes at 37°C. Then, 1X PBS was added to each well to prepare for imaging.

### Propargyl choline labeling of choline-containing phospholipids on fixed cells

Hela cells were cultured in Dulbecco’s modified Eagle’s medium (DMEM) supplemented with 10% FBS for three days, then harvested and resuspended in fresh media. Cells were seeded into an 18-well chamber (Cellvis), 1.5 μl of propargyl choline (10mM; Jena Bioscience) was added to each well, and the total volume per well was 150 μl, and incubated for 24h.

Cells were rinsed with 1X PBS and fixed with 4% formaldehyde for 10 minutes. Cells then were washed with 1X PBS/10% formamide for 30 minutes at 37°C, followed by treatment with 10 μM CalFluor 647 Azide, premixed BTTAA-CuSO_4_ complex ([BTTAA]:[CuSO_4_] = 2:1,[CuSO_4_] = 10 μM) or BTT-DNA ligand ([BTT-DNA] = 20μM) and 2.5 mM freshly prepared sodium ascorbate for 30 minutes at 37°C. After the click reaction, the cells were rewashed with 150 μl wash buffer for 30 minutes twice at 37°C and counterstain with DAPI stain diluted with 1X PBS/10% formamide for 20 minutes at 37°C. Then, 1X PBS was added to each well to prepare for imaging.

### Click extracellular labeling of sialylated glycans on live cells

Hela cells were cultured in Dulbecco’s modified Eagle’s medium (DMEM) supplemented with 10% FBS for three days, then harvested and resuspended in fresh media. 2.5 μl of Ac_4_ManNAl (5mM; Click Chemistry Tools) was added to each well of the 18-well chamber (Cellvis) and dried for 20 min. Cells were seeded into the Ac_4_ManNAl-treated 18-well chamber, and the total volume per well was 150 μl and incubated for 24h.

Cells were washed 3X with PBS/1%FBS (pH 7.4) followed by treatment with 10 μM CalFluor 647 Azide (Click Chemistry Tools), premixed BTTAA-CuSO_4_ complex ([BTTAA]:[CuSO_4_]= 2:1, [CuSO_4_] = 10 μM) or BTT-DNA ligand ([BTT-DNA] = 20 μM) and 2.5 mM freshly prepared sodium ascorbate for 30 minutes on ice. The cells were washed 3X with 200 μl of PBS/1%FBS (pH 7.4) to remove the unreacted CalFluor 647 Azide dye and followed by staining with 100 μl of diluted ReadyProbes™ Cell Viability Imaging Kit, Blue/Green (ThermoFisher Scientific) for 10 minutes at 37°C to stain the nucleus of the live and dead cells. The cells were then washed 3X with PBS/1%FBS (pH 7.4). Then 100 μl of PBS/1%FBS (pH 7.4) was added to each well to prepare for imaging.

### Click extracellular labeling of choline-containing phospholipids on live cells

Hela cells were cultured in Dulbecco’s modified Eagle’s medium (DMEM) supplemented with 10% FBS for three days, then harvested and resuspended in fresh media. Cells were seeded into an 18-well chamber (Cellvis), and 1.5 μl of propargyl choline (10mM; Jena Bioscience) was added to each well, and the total volume per well was 150 μl and incubated for 24h.

Cells were washed 3X with PBS/1%FBS (pH 7.4) followed by treatment with 10 μM CalFluor 647 Azide (Click Chemistry Tools), premixed BTTAA-CuSO_4_ complex ([BTTAA]:[CuSO_4_] = 2:1, [CuSO_4_] = 10 μM) or BTT-DNA ligand ([BTT-DNA] = 20 μM) and 2.5 mM freshly prepared sodium ascorbate for 30 minutes on ice. The cells were washed 3X with 200 μl of PBS/1%FBS (pH 7.4) to remove the unreacted CalFluor 647 Azide dye and followed by staining with 100 μl of diluted ReadyProbes™ Cell Viability Imaging Kit, Blue/Green (ThermoFisher Scientific) for 10 minutes at 37°C to stain the nucleus of the live and dead cells. The cells were then washed 3X with PBS/1%FBS (pH 7.4). Then 100 μl of PBS/1%FBS (pH 7.4) was added to each well to prepare for imaging.

### Click intracellular labeling of choline-containing phospholipids on live cells with lipofectamine RNAiMAX

Hela cells were cultured in Dulbecco’s modified Eagle’s medium (DMEM) supplemented with 10% FBS for three days, then harvested and resuspended in fresh media. Cells were seeded into an 18-well chamber (Cellvis) (2000 cells each well), 1.5 μl of propargyl choline (10mM; Jena Bioscience) was added to each well, and the total volume per well was 150 μl, and incubated for 24h.

12 μl of BTT-DNA ligand (100 μM) and 12 μl of M20-right_3’azide-complementary (100 μM) complementary to BTT-DNA sequence were added to separate 1.5 mL eppendorf™ DNA LoBind microcentrifuge tubes (Fisherscientic) and dried out with SpeedVac SPD1030 integrated vacuum concentrator (Thermo Scientific) at program 1 for 20 minutes. 3 μl of 1XPBS was added to each tube, and we vortexed the tube for 1 minute to dissolve the BTT-DNA and splint. After vortexing, we mixed and hybridized them for 1 hour at 37°C. After hybridization, we diluted the BTT-DNA ligand by adding 15.15 μl prewarmed Opti-MEM™ I reduced serum medium (Thermo Fisher Scientific) to the tube. At the same time, we diluted 0.5 μl of Lipofectamine™ RNAiMAX transfection reagent (Thermo Fisher Scientific) in 15.15 μl prewarmed Opti-MEM™ I reduced serum medium in a different 1.5 ml eppendorf™ DNA LoBind microcentrifuge tube. Then, we added all the diluted BTT-DNA ligands to the Lipofectamine™ RNAiMAX transfection reagent and incubated them at room temperature for 20 minutes to form the DNA-lipid complex. After the encapsulation, 3.2 μl of CF 647 azide (1 mM) was mixed into the DNA-lipid complex. Then, the mixture was applied to cells and incubated for 4 hours in a 5% CO_2_ incubator at 37°C. Cells were then washed 3X with non-phenol red DMEM/10%FBS to remove extra dye and BTT-DNA ligand outside of the cells, followed by treatment with 2 mM freshly prepared sodium ascorbate for 25 minutes in a 5% CO_2_ incubator at 37°C. The cells were washed 3X with 100 μl of non-phenol red DMEM/10%FBS to remove extra sodium ascorbate, followed by staining with 100 μl of diluted ReadyProbes™ Cell Viability Imaging Kit, Blue/Green (Thermo Fisher Scientific) for 5 minutes at 37°C to stain the nucleus of the live and dead cells. The cells were then washed 3X with non-phenol red DMEM/10%FBS. Then 100 μl of non-phenol red DMEM/10%FBS was added to each well to prepare for imaging. For the negative control, untreated cells cultured in propargyl choline and treated cells cultured without propargyl choline were included. Cells treated with BTTAA-CuSO_4_ complex ([BTTAA]:[CuSO_4_]=1:1, [CuSO_4_]= 300 μM) were also included for comparison.

### Click intracellular and extracellular labeling of choline-containing phospholipids on live cells

Hela cells were cultured in Dulbecco’s modified Eagle’s medium (DMEM) supplemented with 10% FBS for three days, then harvested and resuspended in fresh media. Cells were seeded into an 18-well chamber (Cellvis) (2000 cells each well), 1.5 μl of propargyl choline (10mM; Jena Bioscience) was added to each well, and the total volume per well was 150 μl, and incubated for 24h.

12 μl of BTT-DNA ligand (100 μM) and 12 μl of M20-right_3’azide-complementary (100 μM) complementary to BTT-DNA sequence were added to separate 1.5 mL eppendorf™ DNA LoBind microcentrifuge tubes (Fisherscientic) and dried out with SpeedVac SPD1030 integrated vacuum concentrator (Thermo Scientific) at program 1 for 20 minutes. 3 μl of 1XPBS was added to each tube, and we vortexed the tube for 1 minute to dissolve the BTT-DNA and splint. After vortexing, we mixed and hybridized them for 1 hour at 37°C. After hybridization, we diluted the BTT-DNA ligand by adding 15.15 μl prewarmed Opti-MEM™ I reduced serum medium (Thermo Fisher Scientific) to the tube. At the same time, we diluted 0.5 μl of Lipofectamine™ RNAiMAX transfection reagent (Thermo Fisher Scientific) in 15.15 μl prewarmed Opti-MEM™ I reduced serum medium in a different 1.5 ml eppendorf™ DNA LoBind microcentrifuge tube. Then, we added all the diluted BTT-DNA ligands to the Lipofectamine™ RNAiMAX transfection reagent and incubated them at room temperature for 20 minutes to form the DNA-lipid complex. After the encapsulation, 3.2 μl of CF 647 azide (1 mM) was mixed into the DNA-lipid complex. Then, the mixture was applied to cells and incubated for 4 hours in a 5% CO_2_ incubator at 37°C. Cells were then washed 3X with non-phenol red DMEM/10%FBS to remove extra dye and BTT-DNA ligand outside of the cells, followed by treatment with 2 mM freshly prepared sodium ascorbate for 25 minutes in a 5% CO_2_ incubator at 37°C. The cells were washed 3X with 100 μl of non-phenol red DMEM/10%FBS to remove extra sodium ascorbate, followed by 5-minute CuAAC using 10 μM Alexa Fluot™ 555 azide (Thermo Fisher Scientific) in the presence of BTTAA-CuSO_4_ complex ([BTTAA]:[CuSO_4_]=2:1, [CuSO_4_]= 100 μM) and 2 mM sodium ascorbate in a 5% CO_2_ incubator at 37°C. The cells were washed 3X with 100 μl of non-phenol red DMEM/10%FBS to remove CuAAC components, followed by staining with 100 μl of diluted ReadyProbes™ Cell Viability Imaging Kit, Blue/Green (Thermo Fisher Scientific) for 5 minutes at 37°C to stain the nucleus of the live and dead cells. The cells were then washed 3X with non-phenol red DMEM/10%FBS. Then 100 μl of non-phenol red DMEM/10%FBS was added to each well to prepare for imaging.

### Cellular toxicity assay

#### Viability

Hela cells were cultured in Dulbecco’s modified Eagle’s medium (DMEM) supplemented with 10% FBS for three days, then harvested and resuspended in fresh media. Cells were seeded into an 18-well chamber (Cellvis) (2000 cells per well), the total volume per well 150 μl, and incubated for 24h.

12 μl of BTT-DNA ligand (100 μM) and 12 μl of M20-right_3’azide-complementary (100 μM) complementary to BTT-DNA sequence were added to separate 1.5 mL eppendorf™ DNA LoBind microcentrifuge tubes (Fisherscientic) and dried out with SpeedVac SPD1030 integrated vacuum concentrator (Thermo Scientific) at program 1 for 20 minutes. 3 μl of 1XPBS was added to each tube, and we vortexed the tube for 1 minute to dissolve the BTT-DNA and splint. After vortexing, we mixed and hybridized them for 1 hour at 37°C. After hybridization, we diluted the BTT-DNA ligand by adding 16.35 μl prewarmed Opti-MEM™ I reduced serum medium (Thermo Fisher Scientific) to the tube. At the same time, we diluted 0.5 μl of Lipofectamine™ RNAiMAX transfection reagent (Thermo Fisher Scientific) in 16.35 μl prewarmed Opti-MEM™ I reduced serum medium in a different 1.5 ml eppendorf™ DNA LoBind microcentrifuge tube. Then, we added all the diluted BTT-DNA ligands to the Lipofectamine™ RNAiMAX transfection reagent and incubated them at room temperature for 20 minutes to form the DNA-lipid complex. After encapsulation, 0.8 μl of sodium ascorbate (100 mM) was mixed with the DNA-lipid complex. Then, the mixture was applied to cells and incubated for 5 hours in a 5% CO_2_ incubator at 37°C. Cells were then washed 3X with non-phenol red DMEM/10%FBS to remove extra BTT-DNA ligands outside of the cells, followed by staining with 100 μl of diluted ReadyProbes™ Cell Viability Imaging Kit, Blue/Green (ThermoFisher Scientific) for 10 minutes at 37°C to stain the nucleus of the live and dead cells. The cells were then washed 3X with non-phenol red DMEM/10%FBS. Then 100 μl of non-phenol red DMEM/10%FBS was added to each well to prepare for imaging. Untreated cells and cells treated with 2 mM sodium ascorbate were included for negative control. We treated the cells with 300 μM CuSO_4_ in 2 mM sodium ascorbate for positive toxicity control.

#### ROS detection

Hela cells were cultured in Dulbecco’s modified Eagle’s medium (DMEM) supplemented with 10% FBS for three days, then harvested and resuspended in fresh media. Cells were seeded into an 18-well chamber (Cellvis) (2000 cells per well), the total volume per well 150 μl, and incubated for 24h.

12 μl of BTT-DNA ligand (100 μM) and 12 μl of M20-right_3’azide-complementary (100 μM) complementary to BTT-DNA sequence were added to separate 1.5 mL eppendorf™ DNA LoBind microcentrifuge tubes (Fisherscientic) and dried out with SpeedVac SPD1030 integrated vacuum concentrator (Thermo Scientific) at program 1 for 20 minutes. 3 μl of 1XPBS was added to each tube, and we vortexed the tube for 1 minute to dissolve the BTT-DNA and splint. After vortexing, we mixed and hybridized them for 1 hour at 37°C. After hybridization, we diluted the BTT-DNA ligand by adding 16.35 μl prewarmed Opti-MEM™ I reduced serum medium (Thermo Fisher Scientific) to the tube. At the same time, we diluted 0.5 μl of Lipofectamine™ RNAiMAX transfection reagent (Thermo Fisher Scientific) in 16.35 μl prewarmed Opti-MEM™ I reduced serum medium in a different 1.5 ml eppendorf™ DNA LoBind microcentrifuge tube. Then, we added all the diluted BTT-DNA ligands to the Lipofectamine™ RNAiMAX transfection reagent and incubated them at room temperature for 20 minutes to form the DNA-lipid complex. After the encapsulation, 0.8 μl of sodium ascorbate (100 mM) was mixed to the DNA-lipid complex. Then, the mixture was applied to cells and incubated for 5 hours in a 5% CO_2_ incubator at 37°C. Cells were then washed 3X with PBS/1%FBS (pH 7.4) to remove extra BTT-DNA ligands outside of the cells, followed by staining with 100 μl of diluted CellROX™ Green Reagent (Thermo Fisher Scientific) and NucBlue™ Live ReadyProbes™ Reagent (Thermo fisher scientific) for 20 minutes at 37°C. The cells were then washed 3X with PBS/1%FBS (pH 7.4). Then 100 μl of PBS/1%FBS (pH 7.4) was added to each well to prepare for imaging. Untreated cells and cells treated with 2 mM sodium ascorbate were included for negative control. We treated the cells with 300 μM CuSO_4_ in 2 mM sodium ascorbate for positive toxicity control.

#### Mitochondrial Superoxide detection

Hela cells were cultured in Dulbecco’s modified Eagle’s medium (DMEM) supplemented with 10% FBS for three days, then harvested and resuspended in fresh media. Cells were seeded into an 18-well chamber (Cellvis) (2000 cells per well), the total volume per well 150 μl, and incubated for 24h.

12 μl of BTT-DNA ligand (100 μM) and 12 μl of M20-right_3’azide-complementary (100 μM) complementary to BTT-DNA sequence were added to separate 1.5 mL eppendorf™ DNA LoBind microcentrifuge tubes (Fisherscientic) and dried out with SpeedVac SPD1030 integrated vacuum concentrator (Thermo Scientific) at program 1 for 20 minutes. 3 μl of 1XPBS was added to each tube, and we vortexed the tube for 1 minute to dissolve the BTT-DNA and splint. After vortexing, we mixed and hybridized them for 1 hour at 37°C. After hybridization, we diluted the BTT-DNA ligand by adding 16.35 μl prewarmed Opti-MEM™ I reduced serum medium (Thermo Fisher Scientific) to the tube. At the same time, we diluted 0.5 μl of Lipofectamine™ RNAiMAX transfection reagent (Thermo Fisher Scientific) in 16.35 μl prewarmed Opti-MEM™ I reduced serum medium in a different 1.5 ml eppendorf™ DNA LoBind microcentrifuge tube. Then, we added all the diluted BTT-DNA ligands to the Lipofectamine™ RNAiMAX transfection reagent and incubated them at room temperature for 20 minutes to form the DNA-lipid complex. After the encapsulation, 0.8 μl of sodium ascorbate (100 mM) was mixed into the DNA-lipid complex. Then, the mixture was applied to cells and incubated for 5 hours in a 5% CO_2_ incubator at 37°C. Cells were then washed 3X with PBS/1%FBS (pH 7.4) to remove extra BTT-DNA ligands outside of the cells, followed by staining with 100 μl of diluted MitoSOX Green superoxide indicators and NucBlue™ Live ReadyProbes™ Reagent (ThermoFisher Scientific) for 30 minutes at 37°C. The cells were then washed 3X with PBS/1%FBS (pH 7.4). Then 100 μl of PBS/1%FBS (pH 7.4) was added to each well to prepare for imaging. Untreated cells and cells treated with 2 mM sodium ascorbate were included for negative control. We treated the cells with 300 μM CuSO_4_ in 2 mM sodium ascorbate for positive toxicity control.

### Epifluorescent microscopy of cultured cells

For HeLa cells, microscopy was performed using a Nikon inverted research microscope eclipse Ti2-E/Ti2-E/B using a Plan Apo λ 20X/0.75 objective or Plan Apo λ 60X/1.40 oil objective. The Epi-fi LED illuminator linked to the microscope assured illumination and controlled the respective brightness of four types of LEDs of different wavelengths. Images were acquired using the Neutral Density (ND16) filter for labeling with CalFluor 647 Azide. Images were acquired and processed using ImageJ and were shown as a single z-plane. Images acquired using the Neutral Density (ND16) filter are false-colored gray.

## Data Availability Statement

The data supporting this study’s findings are available from the corresponding author upon reasonable request.

## Author Contributions

K.N. and S.H.R. designed the project, analyzed the data, and wrote the manuscript with input from all authors. K.N. performed all the experiments for this project. L.B. helped synthesize the BTT-DNA ligand, and Y.L. helped with the 5-ethynyl uridine (EU) labeling of nascent RNAs in fixed cells using commercial ligands. Y.Q. helped with the copper-induced toxicity assay. Z.Z. helped with the analysis of the copper-induced toxicity assay. M.W. helped with the analysis of fluorogenic plate-reader assay.

