## Supplementary Information for "DNA-enhanced CuAAC ligand enables live-cell detection of intracellular biomolecules"

*Photomicrographs of individual RNA molecule detection with nucleic acid template-driven proximity ligation using BTT-DNA with different CuAAC reaction time*
**Supplementary Figure 8**……10
*Photomicrographs of individual RNA molecule detection with nucleic acid template-driven proximity ligation using BTT-DNA with eight different 5' alkyne DNA linkers*
**Supplementary Figure 9**……11
*Detection and labeling of the intracellular and extracellular biomolecules using CuAAC in the presence of BTT-DNA and $CuSO_4$ and BTTAA on fixed cells.*

*Photomicrographs of biocompatibility of the BTT-DNA ligand to live cells cultured with propargyl choline.*
**Supplementary Figure 16** ……………………………………………………………………….18
*Photomicrographs of propargyl-choline treated live Hela cells in the presence of different CalFluor 647 azide concentrations.*
**Supplementary Figure 17** ……………………………………………………………………….19
*Photomicrographs of the intracellular phospholipids labeling and detection in BTT-DNA presence with different transfection times on live cells.*
**Supplementary Figure 18** ……………………………………………………………………….20
*Photomicrographs of cell viability assay of the intracellular phospholipids labeling and detection in BTT-DNA presence.*
**Supplementary Methods**……………………………………………………………………….21

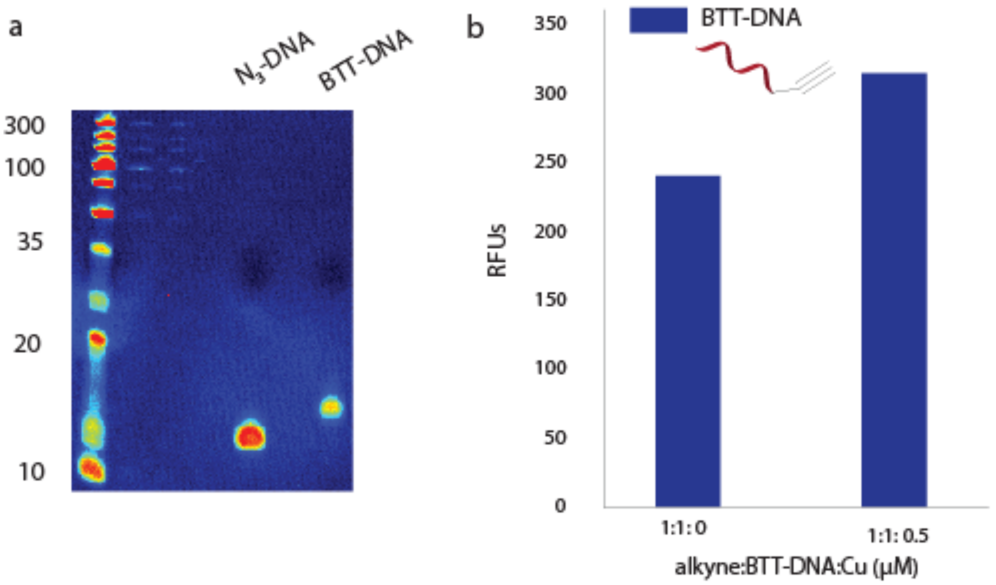

**Supplementary Figure 1.** Characterization of BTT-DNA ligand. a) 15%TBE-Urea gel electrophoresis of BTT-DNA ligand. b) Fluorogenic plate-reader assay showing the ligation of 3-Azido-7-hydroxycoumarin using BTT-DNA at 1 hour.

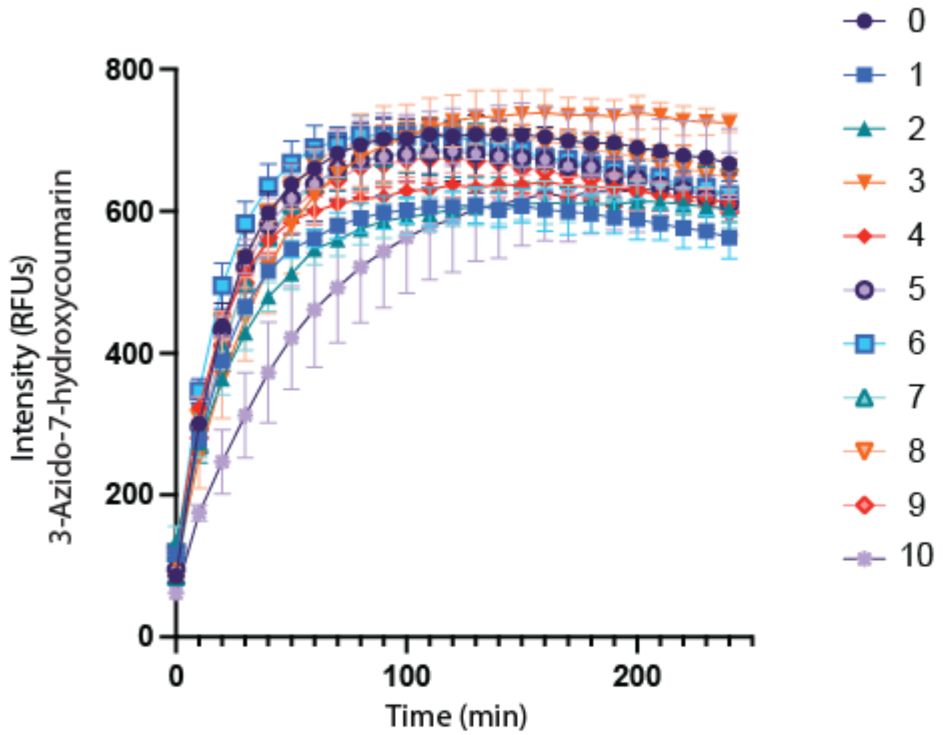

**Supplementary Figure 2.** Stability of BTT-DNA ligand in accelerating CuAAC reaction over ten months. Error bars represent the standard deviation of two biological replicates.

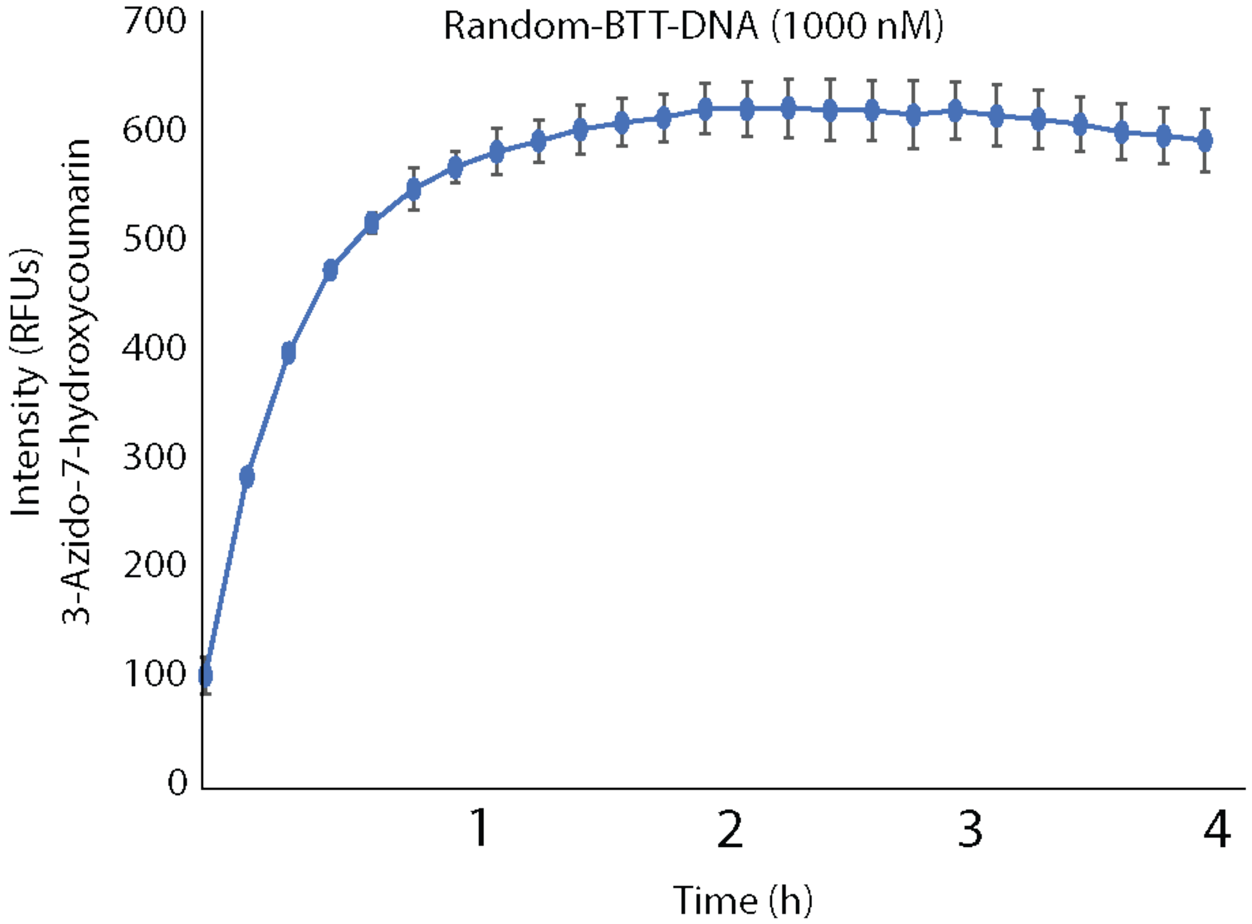

**Supplementary Figure 3.** Reaction kinetics of CuAAC using BTT-DNA with random DNA sequence. Error bars represent the standard deviation of three biological replicates.

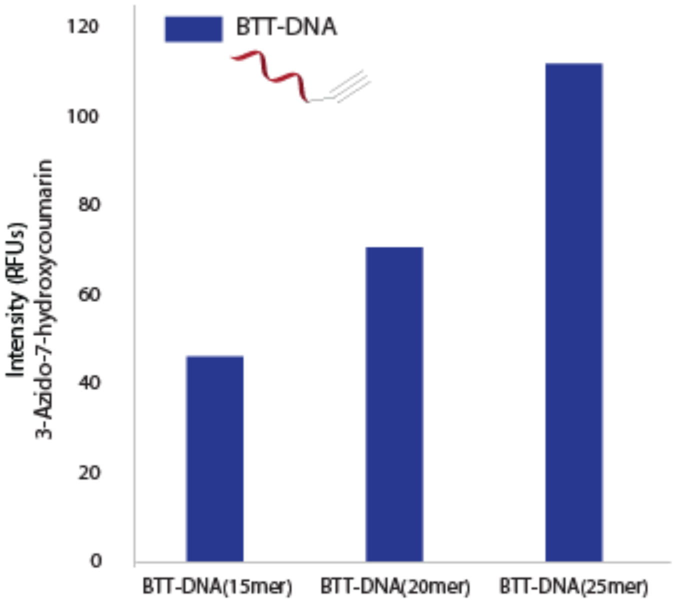

**Supplementary Figure 4.** Fluorogenic plate-reader assay of CuAAC using BTT-DNA ligand with different sequence lengths at 1 hour.

a

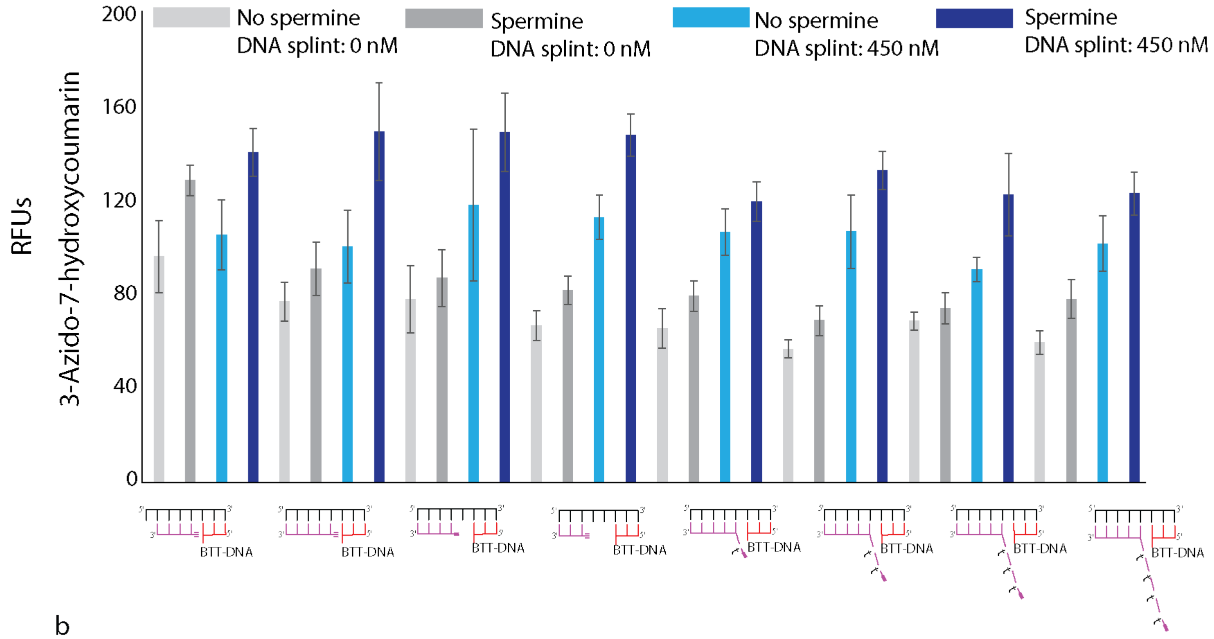

b

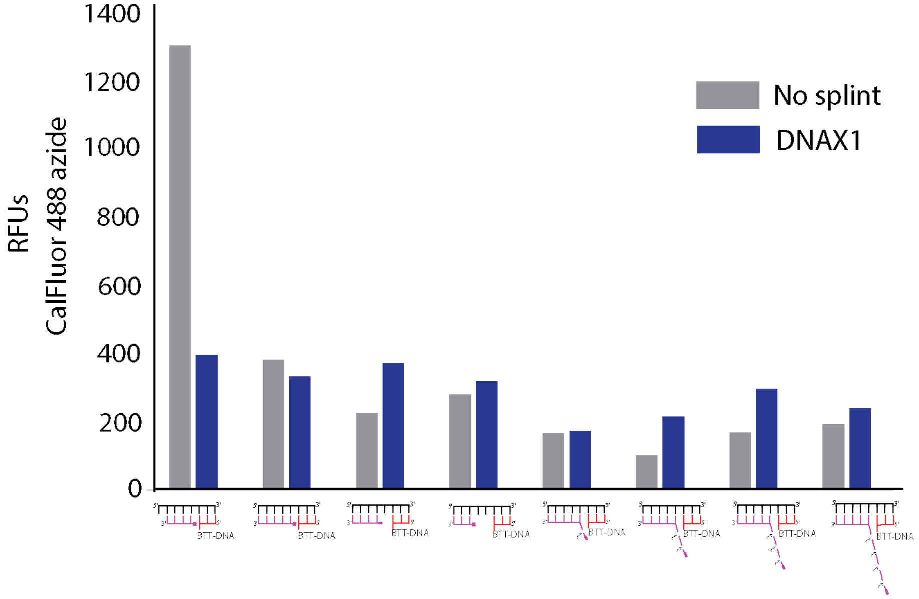

**Supplementary Figure 5.** Optimization of DNA and RNA template-driven proximity ligation with BTT-DNA ligand. a) Comparison of DNA template-driven proximity ligation for 8 different 5' alkyne DNA with/without spermine using 3-Azido-7-hydroxycoumarin. Error bars represent the standard deviation of three biological replicates. b) Comparison of DNA template-driven proximity ligation for 8 different 5' alkyne DNA in the presence of spermine and NaCl using CalFluor 488 azide.

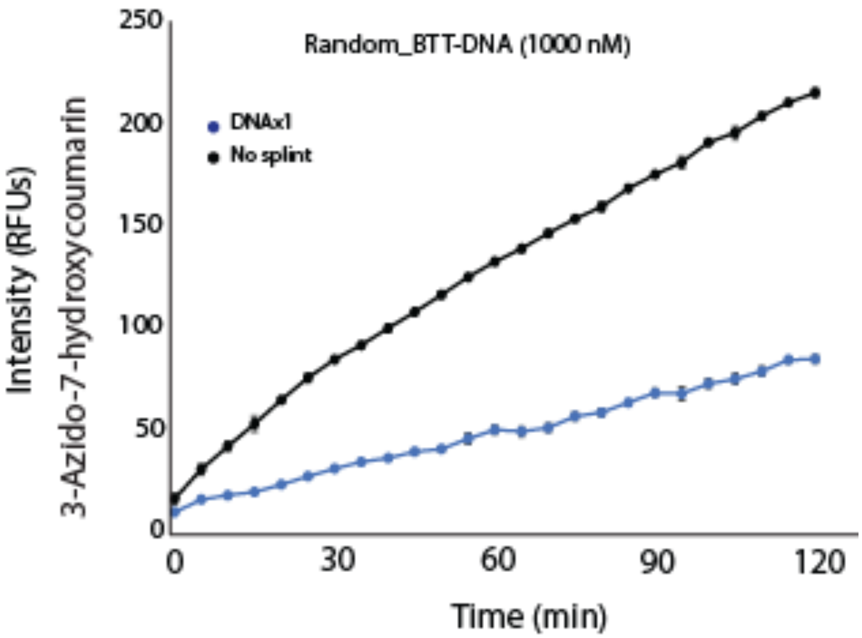

**Supplementary Figure 6.** DNA template-driven proximity ligation using BTT-DNA ligand with random DNA sequence. Each condition has two biological replicates.

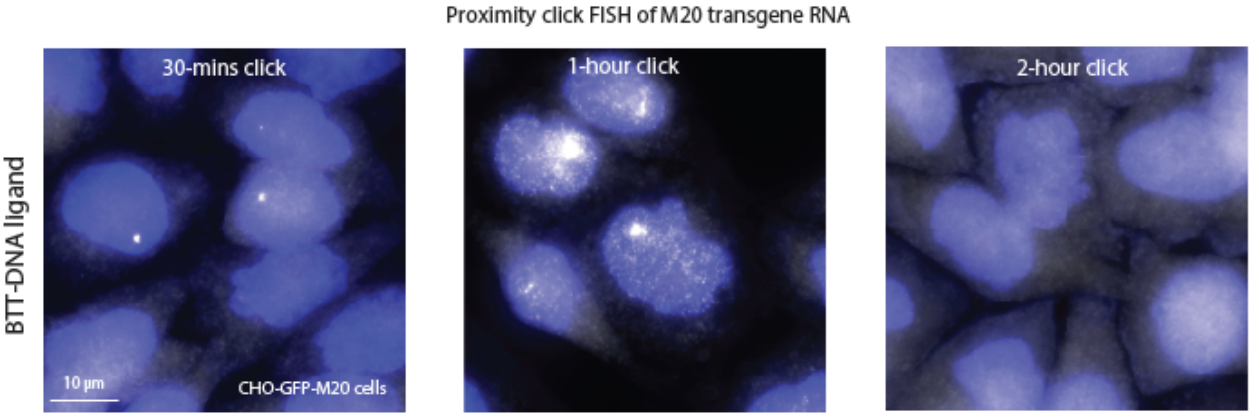

**Supplementary Figure 7.** Photomicrographs of individual RNA molecule detection with nucleic acid template-driven proximity ligation using BTT-DNA with different CuAAC reaction times. Fixed cells are first treated with M20-left-alkyne_20_3t (0.2 µM), then treated with CuAAC in the presence of CalFluor 647 azide (10 µM), BTT-DNA ligand (20 µM), and sodium ascorbate (2.5 mM) for 30 minutes, 1 hour and 2 hours. (white) individual RNA molecule, (blue) DAPI staining of nuclei.

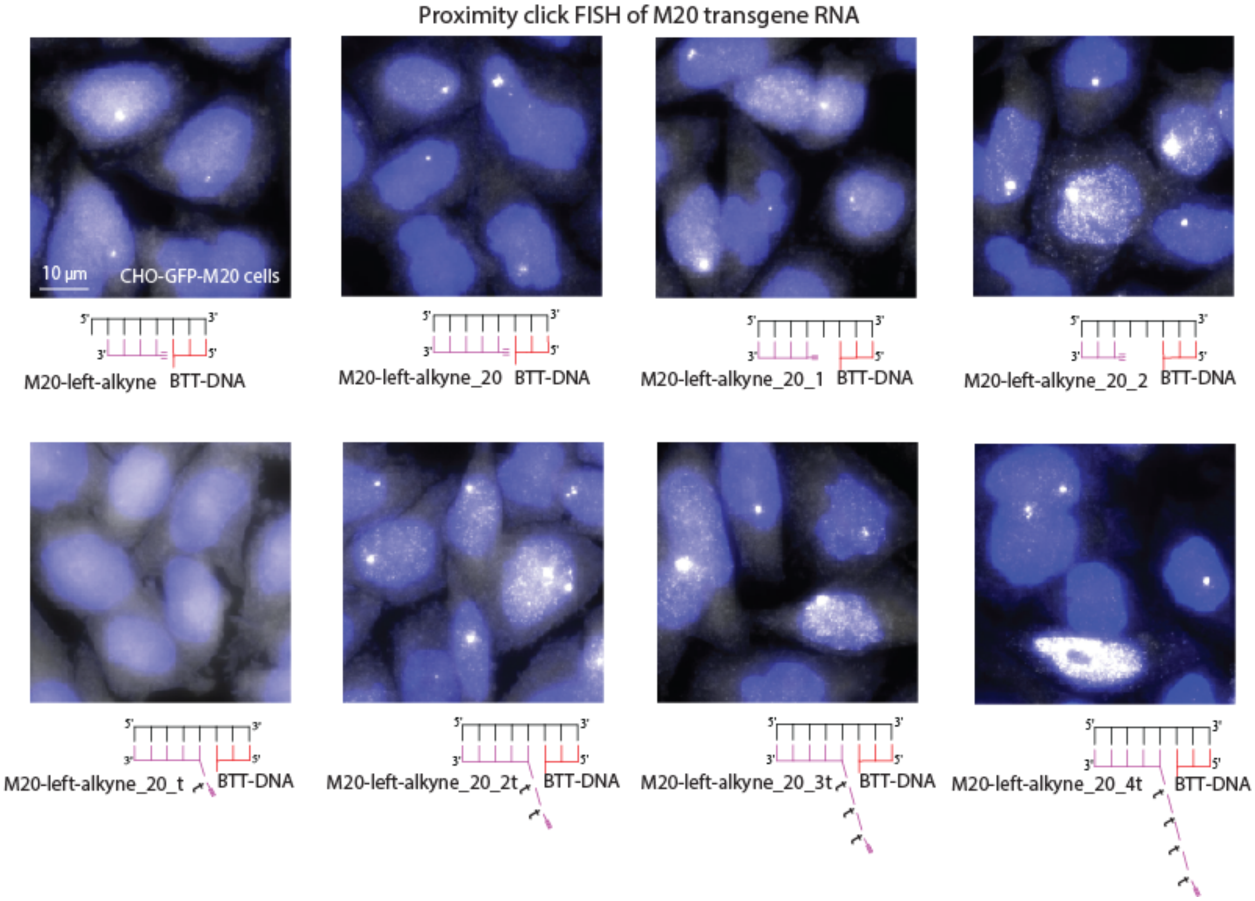

**Supplementary Figure 8.** Photomicrographs of individual RNA molecule detection with nucleic acid template-driven proximity ligation using BTT-DNA with eight different 5’ alkyne DNA. Fixed cells are first treated with 8 different 5’ alkyne DNA (0.2 µM), then treated with CuAAC in the presence of CalFluor 647 azide (10 µM), BTT-DNA ligand (20 µM), and sodium ascorbate (2.5 mM) for 1 hour. (white) individual RNA molecule, (blue) DAPI staining of nuclei.

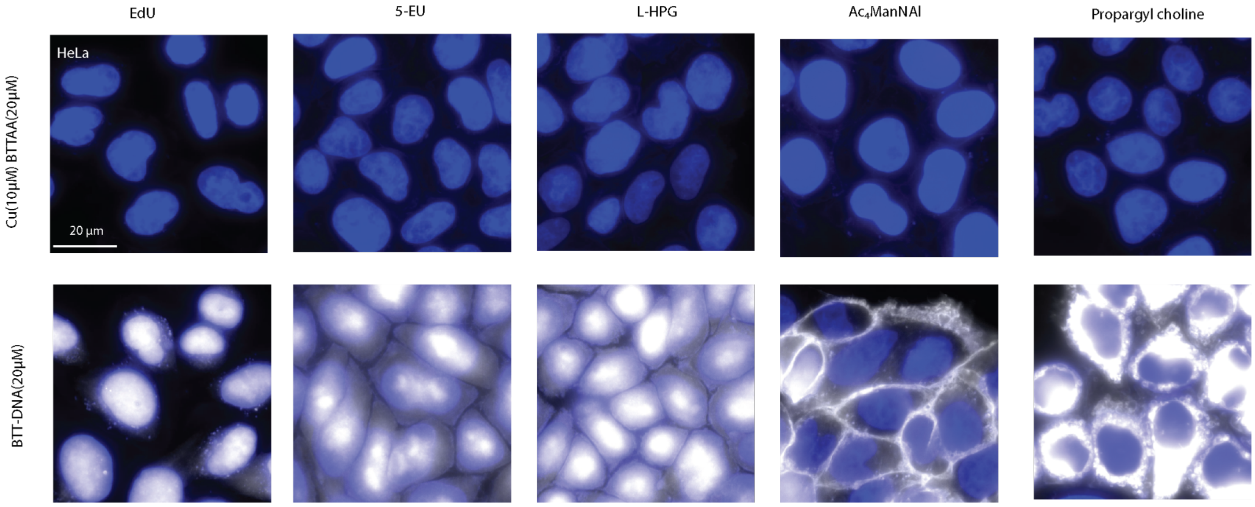

**Supplementary Figure 9.** Detection and labeling of the intracellular and extracellular biomolecules using CuAAC in the presence of BTT-DNA and $CuSO_4$ and BTAA on fixed cells.

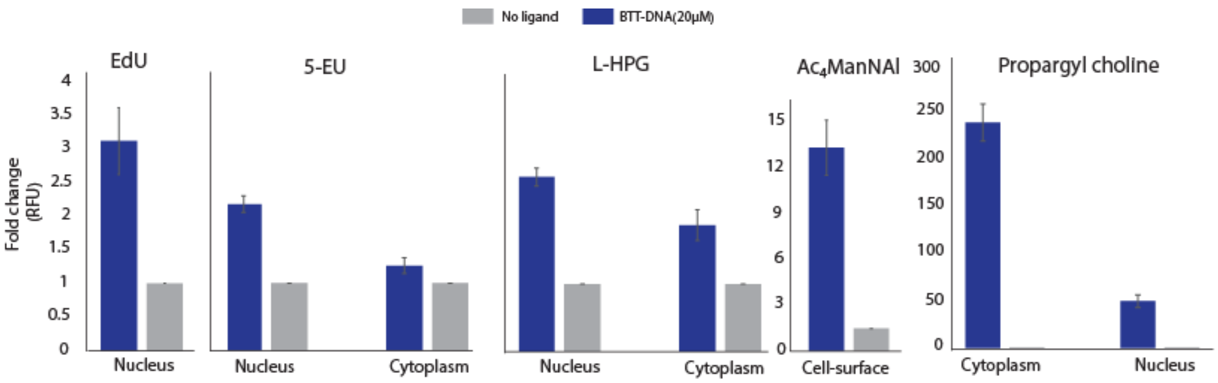

**Supplementary Figure 10.** Quantification of fixed cell detection of biomolecules labeled with CuAAC assisted by BTT-DNA.

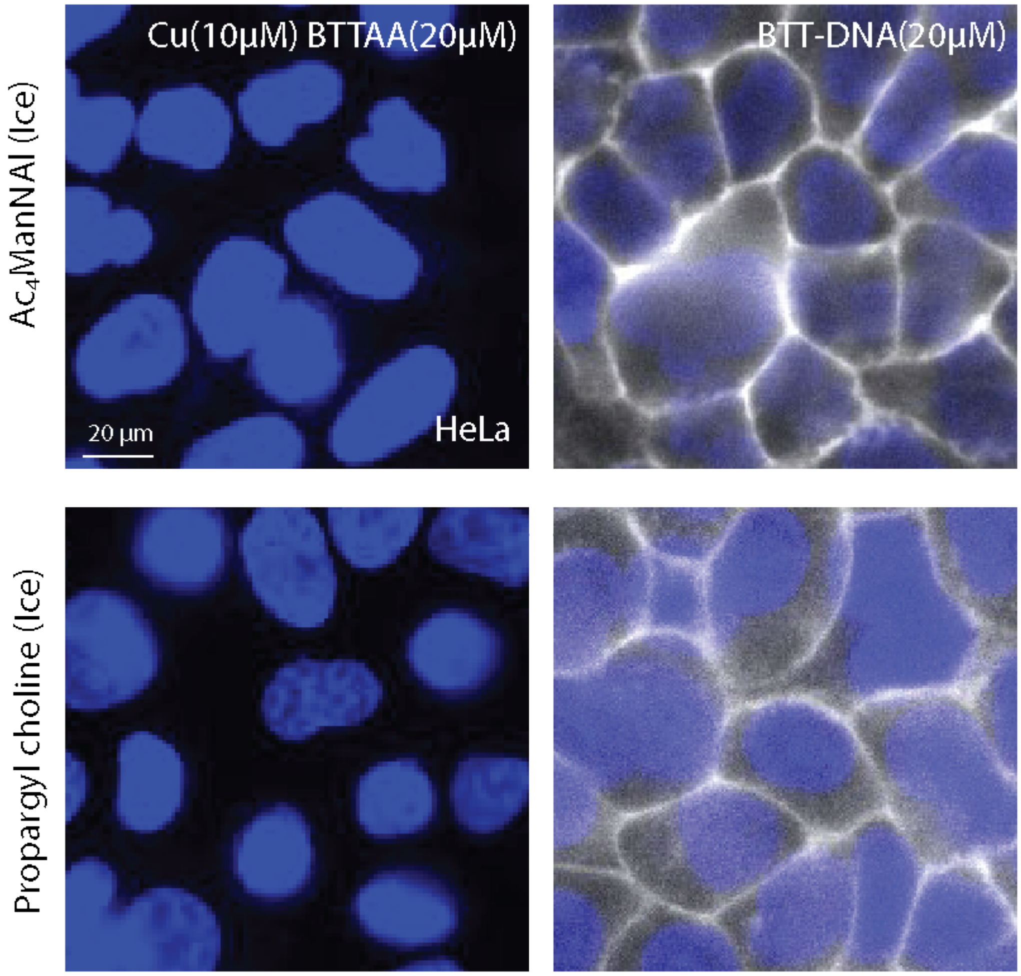

**Supplementary Figure 11.** Detection and labeling of the extracellular biomolecules using CuAAC in the presence of BTT-DNA and $CuSO_4$ and BTAA on live cells.

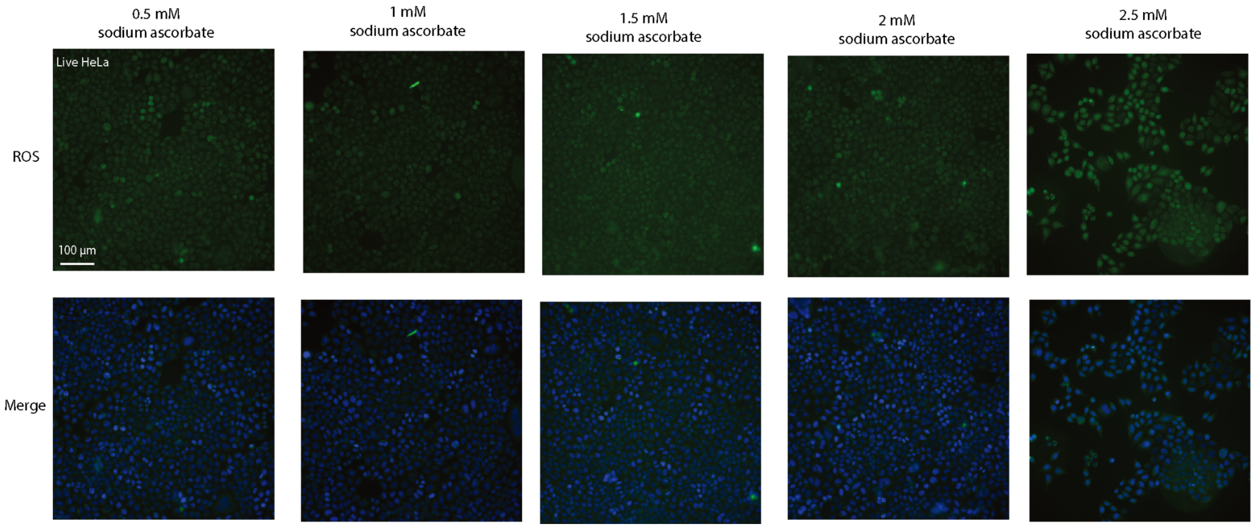

**Supplementary Figure 12.** Representative microscopy of cell reactive oxygen species with different sodium ascorbate concentrations on Hela cells. (blue) Hoechst 33342 staining of nuclei of all cells, (green) CellROX Green Reagent of reactive oxygen species of nuclei of all cells.

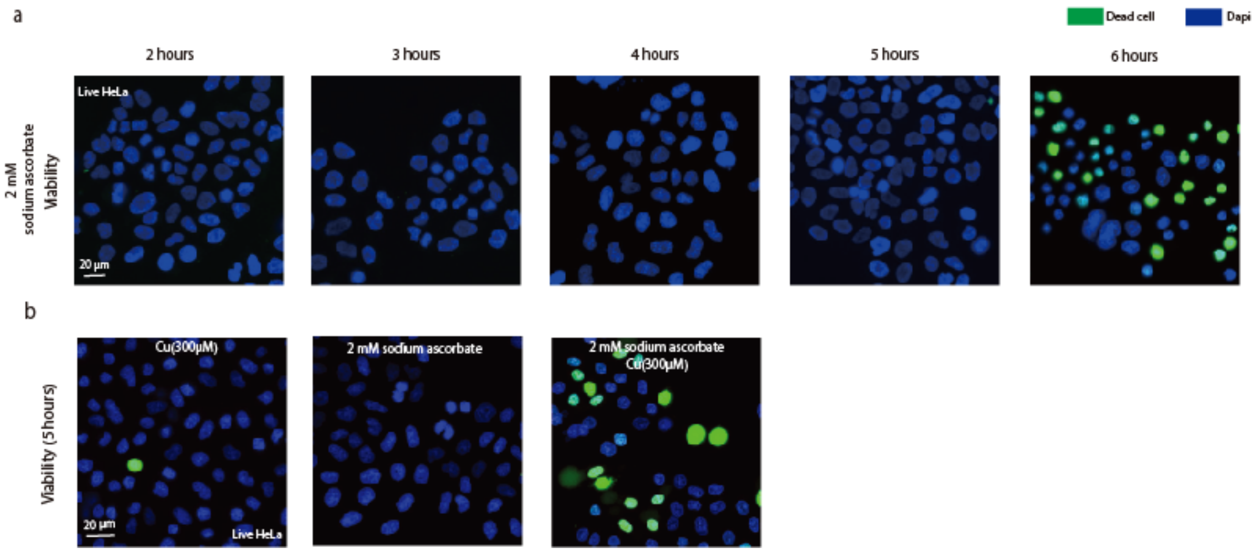

**Supplementary Figure 13.** Representative microscopy of cell viability in the presence of $CuSO_4$ and sodium ascorbate. a) Representative microscopy of cell viability assay with different sodium ascorbate treatment times on Hela cells. b) Comparison of the effect of $CuSO_4$ and sodium ascorbate on cell viability. (blue) Hoechst 33342 staining of nuclei of all cells, (green) SYTOX green nucleic acid staining of nuclei of dead cells.

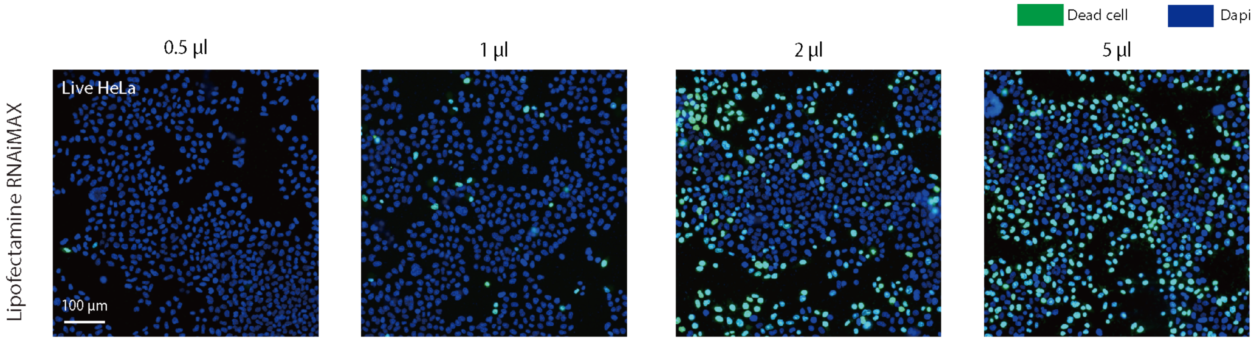

**Supplementary Figure 14.** Photomicrographs of cell viability assay with different lipofectamine RNAiMAX concentrations on Hela cells. (blue) Hoechst 33342 staining of nuclei of all cells, (green) SYTOX green nucleic acid staining of nuclei of dead cells.

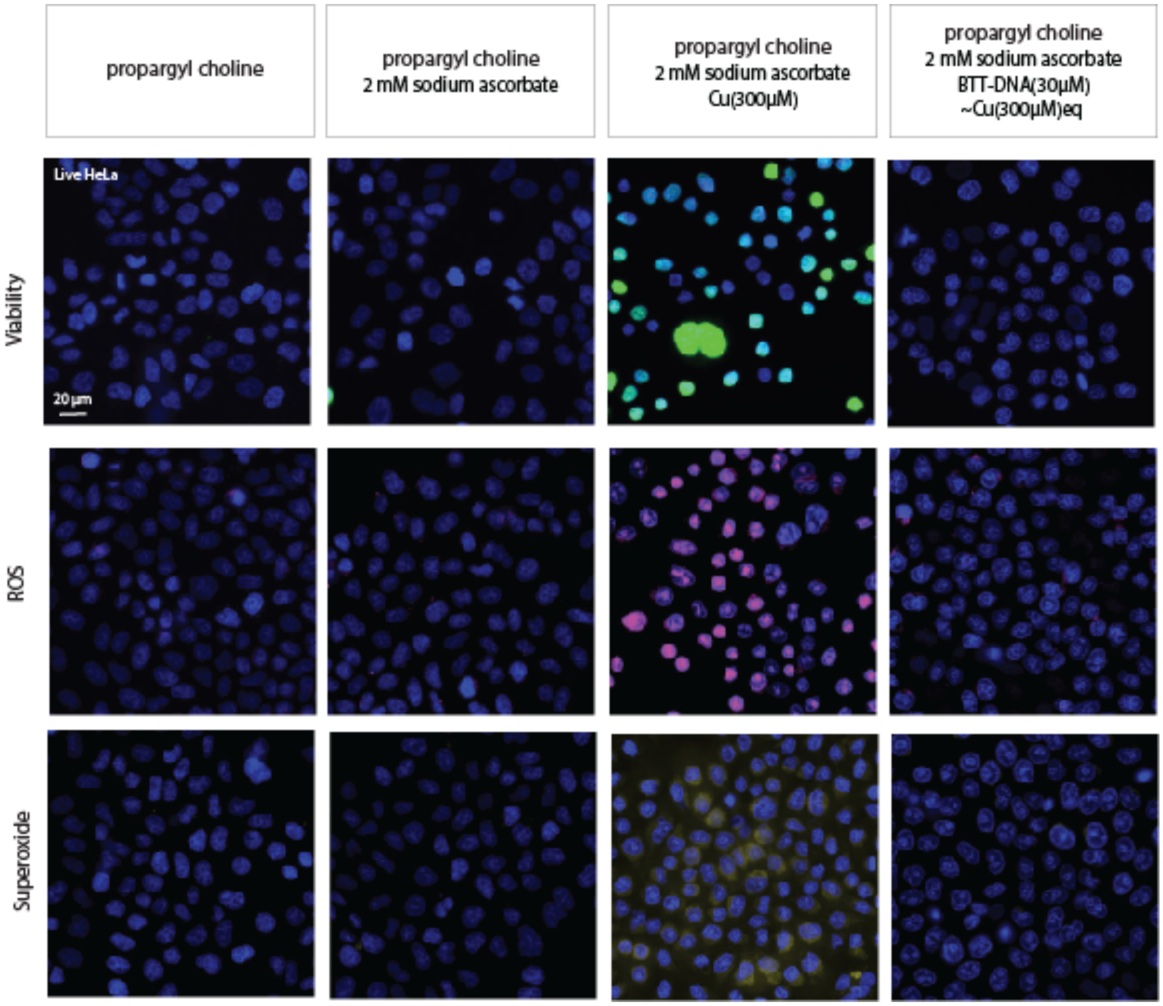

**Supplementary Figure 15.** Photomicrographs of biocompatibility of the BTT-DNA ligand to live cells cultured with propargyl choline.

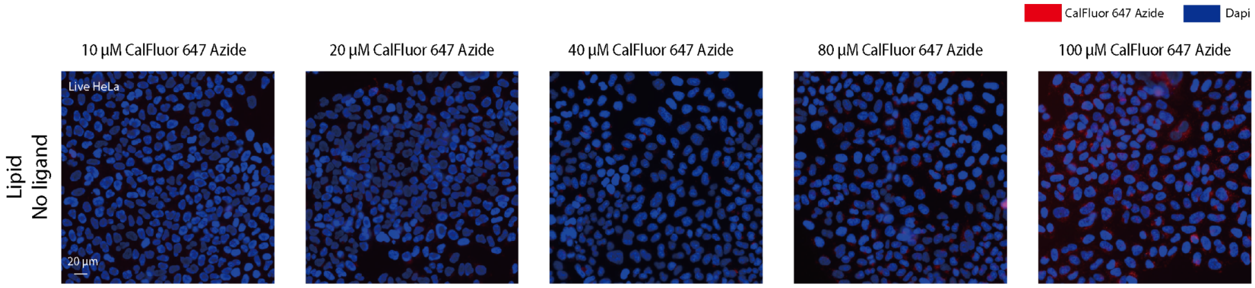

**Supplementary Figure 16.** Photomicrographs of propargyl-choline treated live Hela cells in the presence of different CalFluor 647 Azide concentrations. (red) CalFluor 647 azide, (blue) Hoechst 33342 nuclei staining.

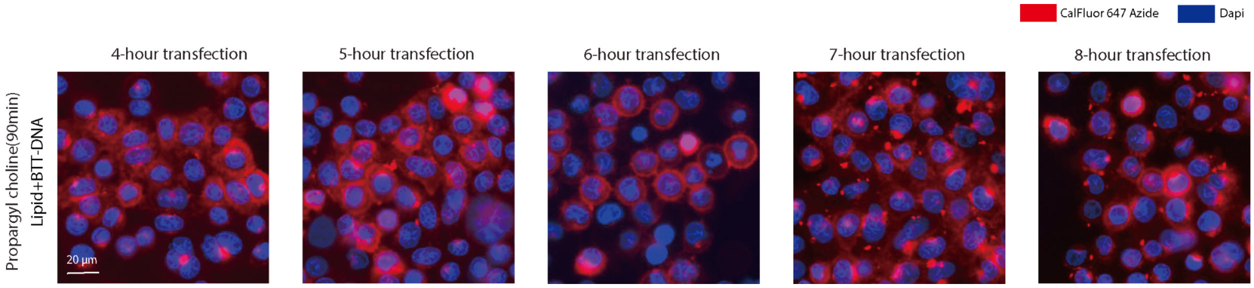

**Supplementary Figure 17.** Photomicrographs of the intracellular phospholipids labeling and detection in BTT-DNA presence with different transfection times on live cells. (red) CalFluor 647 azide labeling of a choline-containing phospholipid, (blue) Hoechst 33342 nuclei staining.

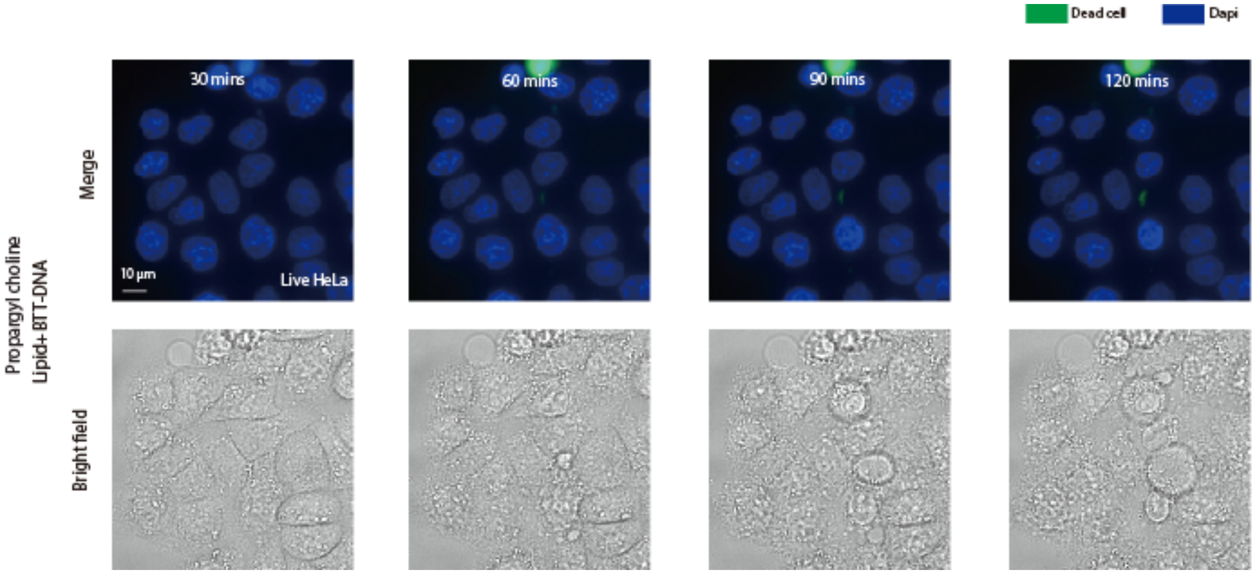

**Supplementary Figure 18.** Photomicrographs of cell viability assay of the intracellular phospholipids labeling and detection in BTT-DNA presence. (blue) Hoechst 33342 staining of nuclei of all cells, (green) SYTOX green nucleic acid staining of nuclei of dead cells.

**Supplementary Methods**

**Synthesis and purification of BTT-DNA ligand.**

A 200-fold molar excess of S1 was reacted with 3’ $N_3$-labeled 15mer ssDNA oligo using BTTAA ligand-assisted CuAAC ([BTTAA]:[$CuSO_4$]=2:1). The reaction mixture was prepared by adding the following reagents: $N_3$-DNA (100 μM), S1-alkyne (40 mM), premixed copper sulfate (5 mM) and BTTAA (20 mM), then sodium ascorbate (100 mM). It was then shaken at 600 rpm for 30 mins at 37°C, resulting in a crude BTT-DNA ligand. The BTT-DNA ligand was purified using 3.5kD MWCO dialysis tubing for 24 hours in 3.5 L of water at room temperature (Repligen). The Invitrogen Qubit ssDNA Assay Kit was used to measure the concentration of the purified BTT-DNA ligand.

| Reagent | stock | Volume added | Final concentration |
|---|---|---|---|
| $N_3$-DNA | 100 μM | 48 μl | 6 μM |
| S1-alkyne | 40 mM | 24 μl | 1200 μM |
| CuSO4 | 5 mM | 12 μl | 75 μM |
| BTTAA | 20 mM | 6 μl | 150 μM |
| Sodium Ascorbate | 100 mM | 20 μl | 2.5 mM |
| NF-H20 | | 690 μl | |
| Total volume | | 800 μl | |

**Click reaction plate assay.**

The click reaction mixture was prepared by adding the following reagents: PBS (10X), alkyne (10 μM), premixed copper sulfate (5 μM), and BTTAA (10 μM) or BTT-DNA(10 μM), azido dye (75 μM), then sodium ascorbate (25 mM). The click reaction mixture was added to the wells (20 μL/well) of the 384-well fluorescence plate, and the click reaction was carried out for 2 hours at room temperature. The fluorescent alkyne-azide cyclo-adduct was detected using Spectral Max Gemini EM plate reader with excitation/emission wavelengths at 404/477 nm for 3-azido-7-hydroxycoumarin, 500/521nm for CalFlour 488 azide, 561/583nm for CalFlour 555 azide, 591/609nm for CalFlour 580 azide, 657/674nm for CalFlour 647 azide. For the DNA and RNA template-driven proximity ligation click reaction, DNA or RNA (4.5 μM), spermine (1 mM), and NaCl (4M) were added after BTT-DNA. Kinetic data was analyzed using Igor Pro software (Wavemetrics)

| Reagent | Stock | Volume added | Final concentration |
|---|---|---|---|
| PBS | 10X | 2 μl | 1X |
| Alkyne | 10 μM | 2 μl | 1 μM |
| BTT-DNA(or Cu and BTTAA premix | 10 μM | 2 μl | 1 μM |

| | | | |
|---|---|---|---|
| Azido dye | 75 µM | 2 µl | 7.5 µM |
| NF-H2O | | 10 µl | |
| Sodium Ascorbate | 25 mM | 2 µl | 2.5 mM |
| Total volume | | 20 µl | |

**Table of the oligonucleotides used.**

| **Oligonucleotide** | **Sequence** |
|---|---|
| M20-right_3'azide | 5'-GGTGCTCTTCGTCCA/3AzideN/ |
| Random-M20_3'azide | 5'-GCA GAG ACA CTA TTG/3AzideN/ |
| M20-left-alkyne | 5Hexynyl/CAAACACAACTCCTG-3' |
| M20-left-alkyne_20 | 5Hexynyl/CAAACACAACTCCTGGTCGA-3' |
| M20-left-alkyne_20_1 | 5Hexynyl/AAACACAACTCCTGGTCGAG-3' |
| M20-left-alkyne_20_2 | 5Hexynyl/AACACAACTCCTGGTCGAGG-3' |
| M20-left-alkyne_20_t | 5Hexynyl/TCAAACACAACTCCTGGTCGA-3' |
| M20-left-alkyne_20_2t | 5Hexynyl/TTCAAACACAACTCCTGGTCGA-3' |
| M20-left-alkyne_20_3t | 5Hexynyl/TTTCAAACACAACTCCTGGTCGA-3' |
| M20-left-alkyne_20_4t | 5Hexynyl/TTTTCAAACACAACTCCTGGTCGA-3' |
| DNAX1 splint | 5'-CGACCTCGACCAGGAGTTGTGTTTGTGGACGAAGAGCACC-3' |
| DNAX2 splint | 5'-CGACCTCGACCAGGAGTTGTGTTTGTGGACGAAGAGCACCAAACGACCTCGACCAGGAGTTGTGTTTGTGGACGAAGAGCACC-3' |
| DNAX3 splint | 5'-CGACCTCGACCAGGAGTTGTGTTTGTGGACGAAGAGCACCAAACGACCTCGACCAGGAGTTGTGTTTGTGGACGAAGAGCACCAAACGACCTCGACCAGGAGTTGTGTTTGTGGACGAAGAGCACC-3' |
| RNAX1 splint | 5'-CGACCUCGACCAGGAGUUGUGUUUGUGGACG |

| | |
|---|---|
| | AAGAGCACC-3' |
| M20-right_3'azide-complementary | 5'-TGGACGAAGAGCACC-3' |

**Quantification of labeling intensity of biomolecules with BTT-DNA-assisted CuAAC on fixed cells.**
Fluorescence intensities were obtained from random points in various raw fluorescence micrographs. A minimum of 175 points were acquired for analysis of EdU labeling, a minimum of 90 points were acquired for EU labeling, a minimum of 115 points were obtained for L-HPG labeling, a minimum of 85 points were acquired for $Ac_4ManNAl$ labeling, and a minimum of 95 points were obtained for propargyl choline labeling. Intensity was normalized to the fluorescence intensity from no ligand control.

**Quantification of extracellular labeling intensity of biomolecules with BTT-DNA-assisted CuAAC on live cells.**
Fluorescence intensities were obtained from random cells in various raw fluorescence micrographs. 5 random points were acquired from each cell membrane to obtain the average fluorescence of each cell. 60 cells were acquired for $Ac_4ManNAl$ labeling, and 45 cells were obtained for propargyl choline labeling. Intensity was normalized to the fluorescence intensity from no ligand control.

**Quantification of intracellular labeling of choline-containing phospholipids on live cells with lipofectamine RNAiMAX.**
Fluorescence intensities were obtained from random 60 cells in three fluorescence micrographs from three biological replicates. 5 random points were acquired from each cell cytoplasm to obtain the average fluorescence of each cell. Intensity was normalized to the fluorescence intensity from the fluorescence micrographs at 30 minutes.

**Analysis for cellular toxicity assay**
**Viability**
The analysis of cell viability was conducted using CellProfiler [1]. The details are in the CellProfiler project file (Viability_analysis.cpproj). Briefly, the Dapi and GFP channels of each image containing many cells were used for analysis. The Dapi channel was used to segment the nuclei by global minimum cross-entropy thresholding on the logarithm of intensity. Each nucleus was counted as one cell. The cells were classified as GFP-positive or GFP-negative based on the mean intensity of the GFP channel in the nucleus, where the threshold was set based on the intensity of the GFP channel from the untreated cells. The intensity of GFP-positive cells is higher than the threshold, and the intensity of GFP-negative cells is lower than the threshold. For the viability analysis, GFP-negative cells are viable cells with intact cell membranes, so they were accepted for further analysis. Sixteen fluorescence microscopy images from two biological experiments were obtained for the ratio of viable cells.

**ROS detection**
The analysis of ROS in cells was conducted using CellProfiler [1]. The details are in the CellProfiler project file (ROS_analysis.cpproj). Briefly, the Dapi and GFP channels of each image containing many cells were used for analysis. The Dapi channel was used to segment

the nuclei by global minimum cross-entropy thresholding on the logarithm of intensity. Each nucleus was counted as one cell. The cells were classified as GFP-positive or GFP-negative based on the mean intensity of the GFP channel in the nucleus, where the threshold was set based on the intensity of the GFP channel from the untreated cells. The intensity of GFP-positive cells is higher than the threshold, and the intensity of GFP-negative cells is lower than the threshold. For the ROS analysis, GFP-positive cells are cells with ROS, so they were accepted for further analysis. Sixteen fluorescence microscopy images from two biological experiments were obtained for the ratio of cells with ROS.

**Mitochondrial Superoxide detection**

For the analysis of mitochondrial superoxide, the threshold for the intensity of mitochondria is based on the intensity of mitochondria from untreated cells because the MitoSOX Green superoxide indicators measure the superoxide produced only by mitochondria. Based on the images, all cells treated with copper in the presence of sodium ascorbate show high fluorescent intensity in the mitochondria compared to the negative controls. The cells treated by BTT-DNA ligand are negative, so we analyzed the positive cells manually rather than using Cell Profiler.

1. Stirling, D. R. *et al.* CellProfiler 4: improvements in speed, utility and usability. *BMC Bioinformatics* **22**, 433 (2021).
